# Single-cell imaging reveals a key role of Bck2 in budding yeast cell size adaptation to nutrient challenges

**DOI:** 10.1101/2024.10.04.616606

**Authors:** Yagya Chadha, Igor V. Kukhtevich, Francesco Padovani, Robert Schneider, Kurt M. Schmoller

## Abstract

Cell size is tightly controlled to optimize cell function and varies broadly depending on the organism, cell type, and environment. The budding yeast *S. cerevisiae* has been successfully used as a model to gain insights into eukaryotic cell size control. Multiple regulators of cell size in steady-state conditions have been identified, such as the G1/S transition activators Cln3 and Bck2 and the inhibitor Whi5. Individual deletions of these regulators result in populations with altered mean cell volumes. However, size homeostasis remains largely intact. Here, we show that although the roles of Bck2 and Cln3 for cell size regulation appear largely redundant in steady-state, a switch from fermentable to non-fermentable growth media reveals a unique role for Bck2 in cell size adaptation to changing nutrients. We use live-cell microscopy and machine learning-assisted image analysis to track single cells and their progeny through the nutrient switch. We find that after the switch, *bck2Δ* cells experience longer cell cycle arrests and more arrest-associated enlargement than wild-type, *whi5Δ* or *cln3Δ* cells, indicating that Bck2 becomes the critical G1/S activator in changing nutrients. Our work demonstrates that studying size regulation during nutrient shifts to mimic the dynamic environments of free-growing microorganisms can resolve apparent redundancies observed in steady-state size regulation.

## Introduction

The size of a cell is a critical regulator of its function and misregulation of cell size is associated with various disease states ^1^. While the underlying causal relationships are often unclear, recent studies have revealed that increased cell size promotes cellular senescence and contributes to aging ^2–4^. It is therefore crucial that cell size is tightly controlled and adapted according to external and internal parameters, such as dynamic environments and developmental changes. Size homeostasis, that is the maintenance of narrow cell size distributions, is attributed to the cell’s ability to measure its size and coordinate cell cycle progression and cell growth with cell size ^5–7^. For example, one common size-sensing strategy that has been reported for budding yeast ^8^, plant ^9^ and mammalian cells ^10,11^ is the cell-growth-based concentration decrease of a cell cycle inhibitor.

However, while such specific size control mechanisms have been identified, there is growing evidence that eukaryotic size homeostasis is a product of multiple size control strategies ^12,13^. In fission yeast, for example, diverse mechanisms integrate cell surface area, cell volume and time information into cell size control ^13^. Similarly, in budding yeast, the concentration of the size-sensing protein Whi5 alone accounts for the size-dependent likelihood of commitment to the G1/S transition in a *bck2Δ* strain ^8^, but multiple other activators and inhibitors of the same cell cycle transition have been identified ^14,15^. Accordingly, disruption of the Whi5-based mechanism only leads to a partial loss of size control ^14,16,17^.

In budding yeast, size control through size-dependent cell cycle progression occurs mainly at the G1/S transition, and dilution by cell growth of the G1/S inhibitor Whi5 is one of the major underlying mechanisms ^8,18^. The G1/S-transition is preceded by the commitment point *Start*, which is largely irreversible due to an underlying positive feedback loop controlling the G1/S transcription factors SBF and MBF (Fig. 1A, ^19,20^). SBF is inhibited by Whi5 and activated by Cln3 and Bck2 ^21–24^. Once the positive feedback loop is triggered, Cln1 and Cln2 are expressed, and in complex with Cdk1, phosphorylate Whi5 to ensure the irreversibility of *Start*. All cells are born with roughly equal amounts of Whi5 and it is diluted by cell growth during G1 ^8,17,18,25–27^, whereas Cln3 and Bck2 concentrations remain mostly constant as G1 progresses ^8,14^. This widening imbalance between SBF activator and inhibitor concentrations is believed to incorporate cell-size information into the cell cycle progression decision ^8,14^ and ultimately trigger the positive feedback loop.

**Figure 1.**
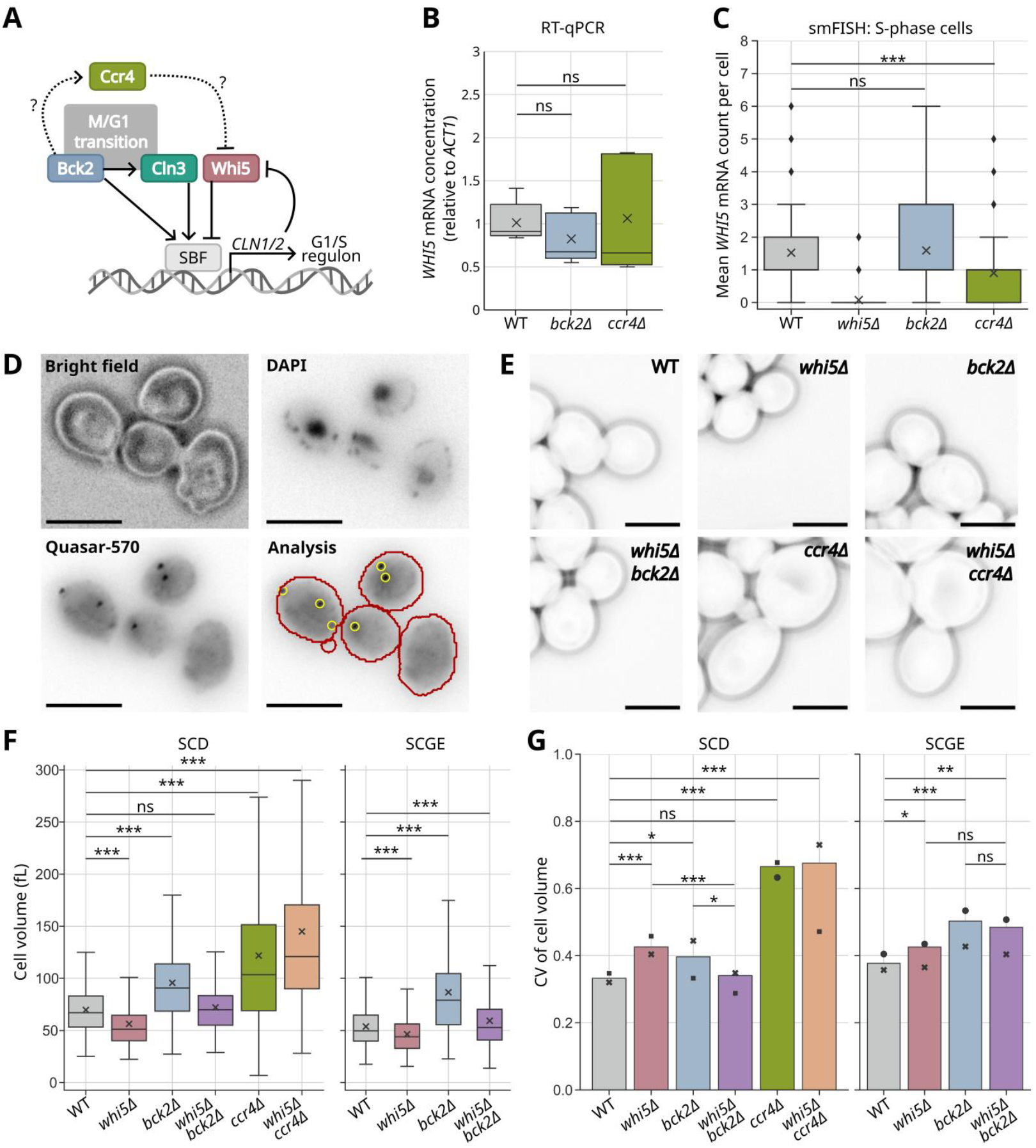
Bck2 acts independently of Whi5. *whi5Δbck2Δ* has wild-type size and more efficient size homeostasis than *whi5Δ* and *bck2Δ* in glucose medium. (**A**) Schematic illustration of key regulators of cell cycle progression and cell size at the budding yeast G1/S transition. (**B**) mRNA concentration of *WHI5* relative to *ACT1* in cells grown in SCD (synthetic complete medium with 2% glucose), as determined from five independent RT-qPCR experiments. x symbol denotes the mean cell volume of the distribution. (**C**) Mean *WHI5* mRNA count per cell from S-phase cells pooled from two independent smFISH experiments performed in SCD (n_WT_ = 172, n_*whi5Δ*_ = 182, n_*bck2Δ*_ = 211, n_*ccr4Δ*_ = 83). x symbol denotes the mean cell volume of the distribution. (**D**) Representative wild type smFISH images: phase contrast; nuclear DNA stained with DAPI; *WHI5* mRNA stained with Quasar-570 labelled smFISH probes; the spots detected using a custom routine in Python in yellow and cell contours in red. Scale bars represent 5 µm. (**E**) Phase-contrast images from steady-state live-cell microscopy in SCD. Representative daughter cells (first-generation cells) from different strains are shown just before division. Scale bars represent 5 µm. (**F**) Cell volume distributions obtained from steady-state live-cell microscopy in SCD and SCGE (synthetic complete medium with 2% glycerol and 1% ethanol). For each strain, all living cells present in the last frame of each imaging position of two independent experiments were pooled. x symbol denotes the mean cell volume of the distribution. Cell numbers (n) for SCD: nWT = 382, n_*whi5Δ*_ = 370, n_*bck2Δ*_ = 376, n_*whi5Δbck2Δ*_ = 463, n_*ccr4Δ*_ = 241, n_*whi5Δccr4Δ*_ = 280. Cell numbers for SCGE: nWT = 533, n_*whi5Δ*_ = 370, n_*bck2Δ*_ = 429, n_*whi5Δbck2Δ*_ = 409. *ccr4Δ* and _*whi5Δccr4Δ*_ cells growing in SCGE were excluded from analysis due to poor growth and high death rate. (**G**) The same dataset was used for calculating the coefficient of variation (CV) of cell volume, as a measure of size homeostasis efficiency. •, ▀, and **x** symbols are used to show the CVs of cell volume calculated from the two independent experimental replicates. For **G**, statistical analyses were performed by comparing overlaps between 10000 bootstrap samples, explained in detail in the Methods. For other subpanels, independent two-tailed t-tests assuming unequal variances (Welch’s t-tests) were used for statistical analyses.

Cln3 and Bck2 are both activators of the G1/S transition. Bck2 activates the G1/S transition in parallel to Cln3 by binding transcription factors Mcm1 and SBF at the *CLN2* promoter ^21,28^. Additionally, it promotes expression of Cln3 and the SBF subunit Swi4 at the M/G1 transition, and of the cyclin Clb2 at the G2/M transition ^21^. Deleting either Bck2 or Cln3 leads to an increased cell volume ^22,29^, and the double deletion causes synthetic lethality ^30^. Despite the changing average cell volumes, size homeostasis, assessed by the coefficient of variation (CV) of cell volume of a population, is surprisingly robust to single deletions of Cln3, Bck2 and Whi5, as well as other *Start* regulators ^14^. While such robustness could be an explanation for why cells evolved multiple redundant *Start* regulators, an alternative explanation is that the apparently redundant regulators fulfil unique roles under specific conditions that have not yet been identified.

Most size control studies so far have been conducted under steady-state conditions, *i*.*e*., during exponential growth in constant environments. Outside laboratory conditions, however, microorganisms frequently encounter rapid changes in the nutritional quality of the environment. Moreover, nutrient availability is known to be a major regulator of cell size, with richer nutrients leading to larger cell sizes ^18,31^. Changing nutrient conditions therefore require dynamic adaptation of cell size, and cell size regulators that appear redundant in constant growth conditions may have specific functions during this size adaptation. Indeed, previous findings point towards specific roles of Cln3 and Bck2 during nutrient changes: the Cln3 protein is very unstable with a half-life on the order of minutes ^32^, and has therefore been proposed to act as a reporter for global cellular protein synthesis ^26,32^. Consistent with such a role, both the Cln3 protein and mRNA amounts are sensitive to nutrient availability ^33,34^, and a switch from a rich to a poor carbon environment is followed by an immediate Cln3 depletion ^26^. Bck2, on the other hand, has been shown to interact with multiple proteins with documented roles in nutrient-sensing pathways ^21^. It has been proposed to be a link between the nutrient status of the environment and passage through the M/G1 transition ^21^.

Here, we use the budding yeast *Saccharomyces cerevisiae* as a model to investigate specific functions of apparently redundant cell size regulators during dynamic cell size adaptation to environmental changes. With microfluidics-based live-cell microscopy and state-of-the-art deep-learning image analysis, we follow the fates of single cells and their complete lineages for multiple generations before and after a nutrient switch from fermentable to non-fermentable carbon sources. By following individual cells, we gain unprecedented insights into the heterogeneity of cell size adaptation and find that the cell cycle state of a cell at the time of the nutrient shift plays a crucial role for the size evolution of its offspring. Moreover, through analysis of deletion mutants, we uncover a unique role for Bck2. We find that after the nutrient switch, size homeostasis is temporarily disrupted, and this disruption is stronger in *bck2Δ* mutants, which show elongated cell cycle arrests. Based on our findings, we propose that Bck2 acts as an essential G1/S transition activator for the period of time following a nutrient switch when Cln3 is depleted. More generally, our work suggests that the apparent redundancy of cell size regulators observed across eukaryotes in steady-state conditions may point to specific roles during dynamic cell size adaptation.

## Results

### Bck2 acts independently of Whi5

Bck2 is mostly thought to promote the G1/S transition independently of Cln3 and Whi5 ^21,28^. However, Manukyan *et al*. ^35^ proposed that Bck2 stimulates the cytoplasmic deadenylase Ccr4, promoting degradation of *WHI5* mRNA (Fig. 1A). Specifically, they found that *WHI5* mRNA was more stable in the absence of Ccr4 in a *GALpr*-*WHI5* strain after switching from galactose to glucose medium. Thus, as a first step to better understand the apparent redundancies of Bck2 and Cln3 for budding yeast G1/S size control, we decided to test this proposed interaction between Bck2, Ccr4 and Whi5 under steady-state conditions and for endogenous Whi5. We used RT-qPCR (Fig. 1B) and single-molecule fluorescence in situ hybridization (smFISH) (Fig. 1C, D, Fig. S1) to measure the effect of deleting *BCK2* or *CCR4* on *WHI5* transcript abundance for cells growing on glucose medium (SCD), but did not see a significant increase. Next, to gain further insights into the genetic interaction of *CCR4* and *WHI5*, we performed time-lapse live-cell microscopy experiments (Fig. 1E-G) and coulter-counter measurements (Fig. S2) of cell volume for single and double deletion strains. We found that *whi5Δ* cells were smaller, and *ccr4Δ* cells were much larger than wild-type cells. However, *whi5Δccr4Δ* did not rescue the large-cell phenotype of *ccr4Δ*. Taken together, our findings are not consistent with Ccr4 impacting cell size through destabilising Whi5 mRNA. Instead, our data shows that Whi5 and Ccr4 affect cell size via independent pathways.

We also performed time-lapse microscopy experiments for *WHI5* and *BCK2* single and double mutants growing on glucose medium (SCD) as well as glycerol-ethanol medium (SCGE). Consistent with previous studies ^14,22,23,36^, *whi5Δ* and *bck2Δ* led to a significant decrease and increase of cell volume, respectively, and the volume of *whi5Δbck2Δ* cells was strikingly similar to that of wild type in both growth media (Fig. 1F, Fig. S3). Thus, the effect of Bck2 and Whi5 on cell size appears to be additive, indicating that Whi5 is not downstream of Bck2 and that they affect cell size via independent paths.

### *whi5Δbck2Δ* has wild-type size and more efficient size homeostasis than *whi5Δ* and *bck2Δ* in glucose medium

After analysing mean cell volumes, we next asked how size control is affected in each of the mutants. The coefficient of variation (CV) of cell volume is the mean-normalised standard deviation of the cell volume distribution, making it a measure of size homeostasis efficiency which is comparable between differently-sized strains. Consistent with their roles in G1/S size control, we found that deletion of either *WHI5* or *BCK2* leads to an increased CV of cell volume, both for cells growing on SCD or SCGE medium (Fig. 1G, Fig. S3). We expected that deletion of both genes would lead to a further reduced size homeostasis efficiency. Surprisingly, the CV of cell volume of *whi5Δbck2Δ* cells was not higher than those of the single deletion mutants (Fig. 1G). For cells growing in SCD, *whi5Δbck2Δ* even showed a significantly decreased CV compared to both Δ*whi5* and Δ*bck2* (Fig. 1G), similar to wild-type cells. Thus, we found that size homeostasis as evaluated by the CV of the cell volume across a steady-state population was surprisingly robust to the double deletion of *WHI5* and *BCK2*, in particular for cells grown in SCD. By contrast, deletion of *CCR4* led to a dramatically increased CV of cell volume for cells grown on glucose, which was not rescued by an additional deletion of *WHI5* (Fig. 1G). This indicates a rare occasion of a drastically weakened size-homeostasis mechanism. However, given that *Δccr4* cells grow very poorly, especially in SCGE, it is unlikely that this lower size homeostasis is a result of disrupted G1/S size control alone.

### The increase in CV of cell volume observed after a nutrient switch is stronger in *whi5Δbck2Δ* than wild type

Prompted by the fact that the absence of Whi5 and Bck2, two major cell size regulators acting at the G1/S transition, does not have a stronger size homeostasis phenotype, even though deletion of each protein individually leads to altered cell volume and weakened size control, we speculated that double deletion of *WHI5* and *BCK2* may impair cell size adaptation upon changing nutrient conditions. To investigate this, we performed a bulk nutrient switch experiment, shifting cells from SCD to SCGE medium (Fig. 2A). Specifically, steady-state populations of cells growing in SCD were washed and inoculated into SCGE (Fig. 2A) at a starting OD_600nm_ of 0.01. We then performed a time course of OD_600nm_ and coulter-counter measurements of cell volume from the time of the nutrient switch up until a new steady-state in the glycerol-ethanol medium was reached (Fig 2B-D). The OD_600nm_ was maintained between 0.1 and 1 through appropriate dilutions. For the first 15 hours after the nutrient switch, both mean cell volume (Fig. 2B) and CV of cell volume (Fig. 2D) increased. Considering the direction of the switch was from a rich medium to a poorer medium, ultimately leading to a smaller mean cell volume post-switch, the initial increase in cell volume was unexpected. This initial increase in mean cell volume and CV of cell volume was not accompanied by notable cell proliferation. Only about 15 hours after the switch, OD_600nm_ started to increase again, and at the same time, mean cell volume and CV of cell volume started to decrease (Fig. 2B, D). Interestingly, during the time of maximal cell volume around 10-20 hours post-switch, *whi5Δbck2Δ* cells showed significant differences compared to wild type. We found that the CV of cell volume of *whi5Δbck2Δ* was significantly higher than that of wild type between 10 and 20 hours post-switch (Fig. 2D). After 20 hours, while the CV of *whi5Δbck2Δ* still appeared to be slightly higher than that of wild type for the remainder of the experiment, the difference was not statistically significant. Thus, both wild-type and *whi5Δbck2Δ cells* showed an increase in cell volume and CV of cell volume post-switch, but *whi5Δbck2Δ* had lower size homeostasis efficiency than wild-type cells. This suggests that the simultaneous loss of both Whi5 and Bck2 does affect size homeostasis but this effect is more prominent under changing nutrient conditions.

**Figure 2.**
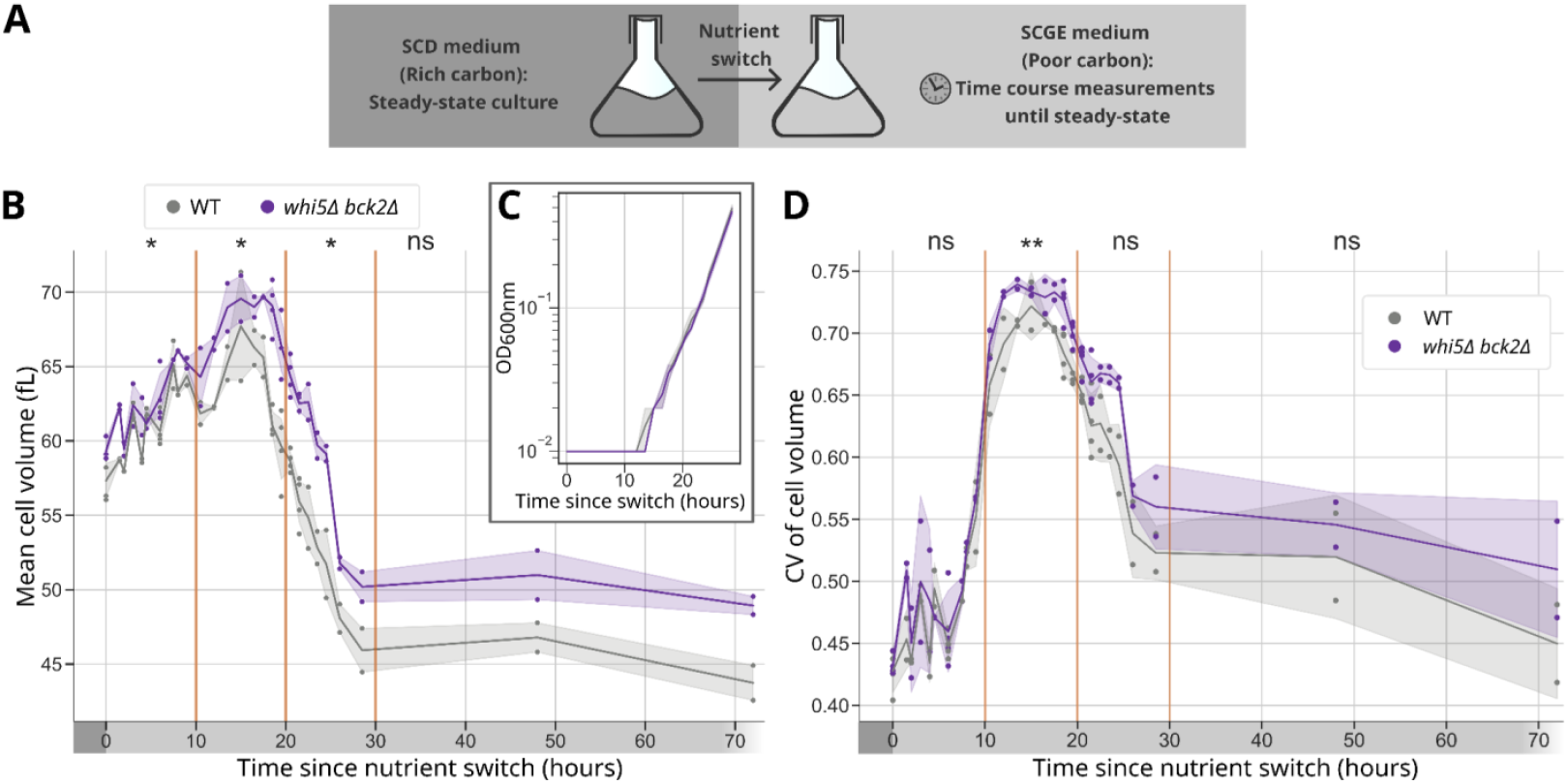
The increase in CV of cell volume observed after a nutrient switch is stronger in *whi5Δbck2Δ* than wild-type cells. (**A**) Illustration of the experiment design for the bulk nutrient switch experiment. Steady-state cells growing in SCD were washed and inoculated into SCGE and a time course of OD600nm and coulter-counter measurements was performed until cells reached steady-state in SCGE. For the measurements taken 48 and 72 hours after the nutrient switch, appropriate dilutions were made when necessary to maintain the OD600nm of the cultures between 0.1 and 1. (**B**) Mean cell volume and OD600nm (**C**, inset) are plotted against time since the nutrient switch. Shaded areas show 95% confidence intervals. Data for each time point are based on at least two experiments. (**D**) The CV of cell volume is plotted against time since the nutrient switch. Shaded areas show 95% confidence intervals. For statistical analysis in **B** and **D**, the time course was divided into three ten-hour windows (orange lines) and the remaining two time points were grouped together and analysed with a mixed ANOVA test (see Methods for details).

### Analysis strategy for live-cell microscopy coupled to nutrient switch

To better understand why both strains underwent an increase in cell volume and CV of cell volume post-switch, and why *whi5Δbck2Δ* cells had a higher CV of cell volume, we performed live single-cell microscopy of cells experiencing the nutrient switch. We grew steady-state cultures of wild type, *whi5Δ, bck2Δ* and *whi5Δbck2Δ* in SCD and transferred them to a custom microfluidic device ^37^ in a time-lapse microscopy setup (Fig. 3A). We imaged the cells at three-minute intervals as they grew in SCD for 2 hours and then automatically switched the medium to SCGE for the next 25 hours. The videos of cell growth generated from this experiment were analysed using the Cell-ACDC pipeline ^38^ and YeaZ ^39^ for cell segmentation (Fig. 3A). The cell masks and tracking were then manually corrected and pedigrees were assigned in a semi-automated fashion in the Cell-ACDC GUI, resulting in a fully-annotated manually corrected dataset of several thousand complete cell cycles (Fig. 3A).

**Figure 3.**
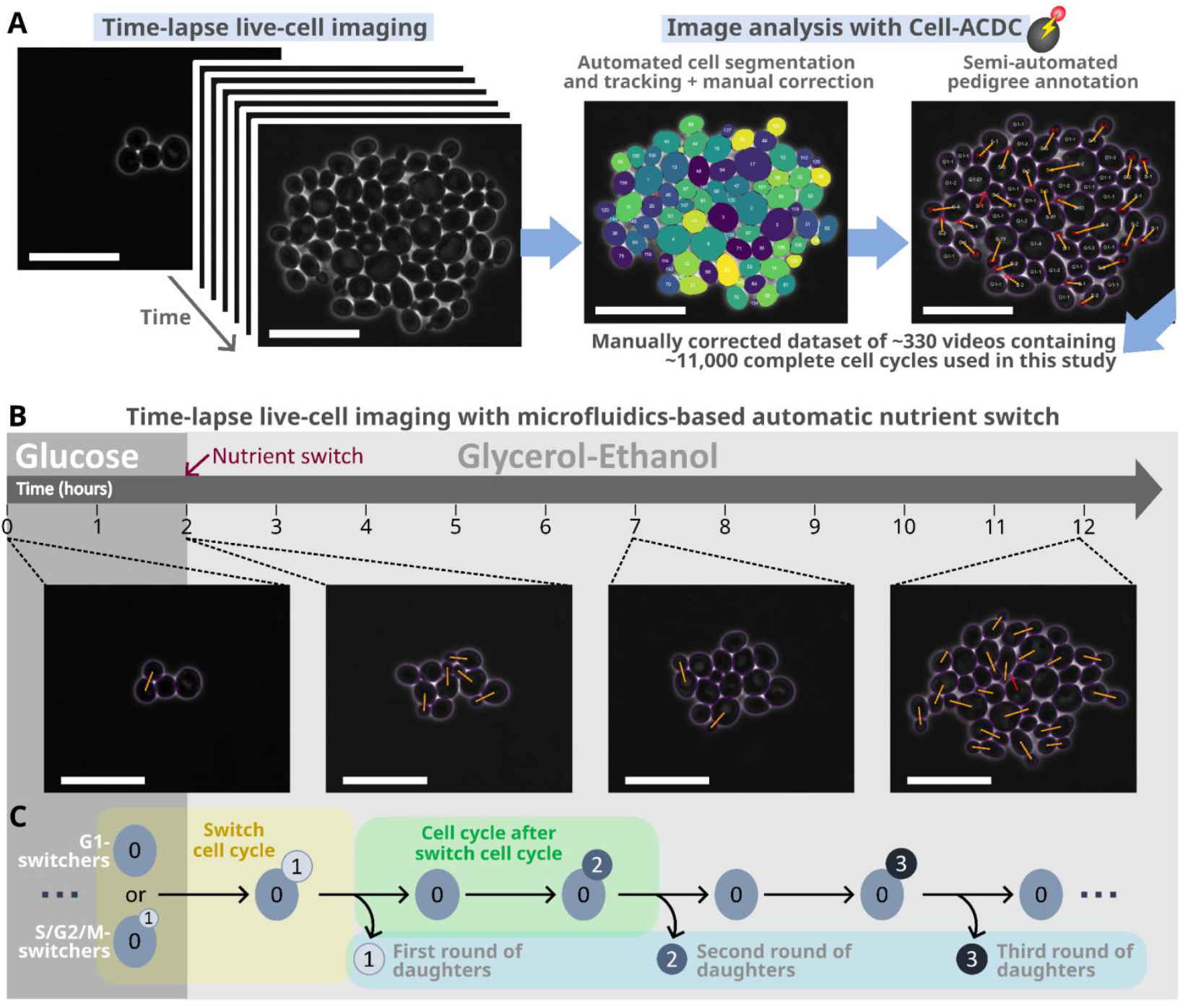
Analysis strategy for live-cell microscopy of the adaptation to the nutrient switch. (**A**) Live-cell imaging analysis pipeline for steady-state and nutrient switch experiments. (**B**) Representative images of WT cells in the nutrient switch live-cell microscopy experiment. (**C**) Cell categories of interest in the downstream analysis of the nutrient switch live-cell microscopy dataset. Scale bars represent 20 µm.

On qualitative inspection of colony growth in the live-cell microscopy videos obtained during the nutrient switch, we found that cells roughly doubled in number during the first two hours in SCD, as is expected from steady-state cells in glucose (Fig. 3B). After the nutrient switch to SCGE, cells arrested for a period of around five hours (WT) but continued to grow in volume. After five hours post-switch, the cells very slowly resumed cell cycle progression. At around 10 hours post-switch, the cell population appeared to have heterogeneous cell sizes, with large cells that had faced the arrest and small cells born after the arrest. These observations are consistent with the increase in the CV of cell volume we observed in the bulk nutrient switch experiment (Fig. 2D). To quantitatively analyse the post-switch adaptation and its effect on cell volume, it is necessary to consider the cell cycle history of each cell. Accordingly, cells were split into multiple categories (Fig. 3C), which we then studied separately.

All cells that were growing in SCD and faced the nutrient switch to SCGE were categorised as ‘*switchers’* (Fig. 3C). At the time of the nutrient switch, the *switchers* could either have been unbudded or budded and were respectively further categorised into ‘*G1-switchers’* or ‘*S/G2/M-switchers’*. The specific cell cycle of the *switchers* during which the nutrient switch occurred was called the *‘switch-cell-cycle’* (Fig. 3C, highlighted in yellow). As these *switchers* overcame the post-switch arrest and finished the *switch-cell-cycle*, they divided to give rise to the *first round of daughters*. The *switchers* then entered the *cell-cycle-after-switch-cell-cycle* (Fig. 3C, highlighted in green), at the end of which they gave rise to the *second round of daughters*. The following rounds of daughters were categorised analogously (Fig. 3C, highlighted in blue). The rounds of daughters could be further categorised into *daughters of G1-switchers* or *daughters of S/G2/M switchers*. Notably, the *first round of daughters of S/G2/M-switchers* was the only category apart from the *switchers* themselves that faced the nutrient switch – as buds of the *S/G2/M-switchers*. Albeit complex, this careful categorisation of cells was key to identifying specific phenotypes in response to the nutrient switch in the following analysis.

### The nutrient switch causes cell cycle arrests in *switchers* and leads to stronger cell enlargement in *bck2Δ* cells

As described in Fig. 4, we analysed two specific cell cycles of *switchers*: the *switch-cell-cycle* (Fig. 4A, highlighted in yellow) and the *cell-cycle-after-switch-cell-cycle* (Fig. 4A, highlighted in green). *G1-switchers* arrested in G1 in the *switch-cell-cycle* (Fig. 4B, left yellow). The following S/G2/M phase of the switch cycle was not strongly affected in length and was similar to that of cells growing exponentially in SCGE. Also the phase lengths in the next cell cycle of *G1-switchers* (*cell-cycle-after-switch-cell-cycle*, Fig. 4B, left green) were similar to those of cells growing exponentially in SCGE. On the other hand, *S/G2/M-switchers* naturally had a normal steady-state G1 length in SCD medium and arrested in S/G2/M post-switch (Fig. 4B, right yellow). After completing the *switch-cell-cycle*, they also exhibited a strongly prolonged G1 phase in the next cycle (Fig. 4B, right green). By contrast, the duration of the subsequent S/G2/M was close to that of cells exponentially growing in SCGE (Fig. 4B, green). These results showed that all cells that faced the nutrient switch immediately arrested in the ongoing cell cycle stage. For *S/G2/M-switchers*, the next G1 was also elongated.

**Figure 4.**
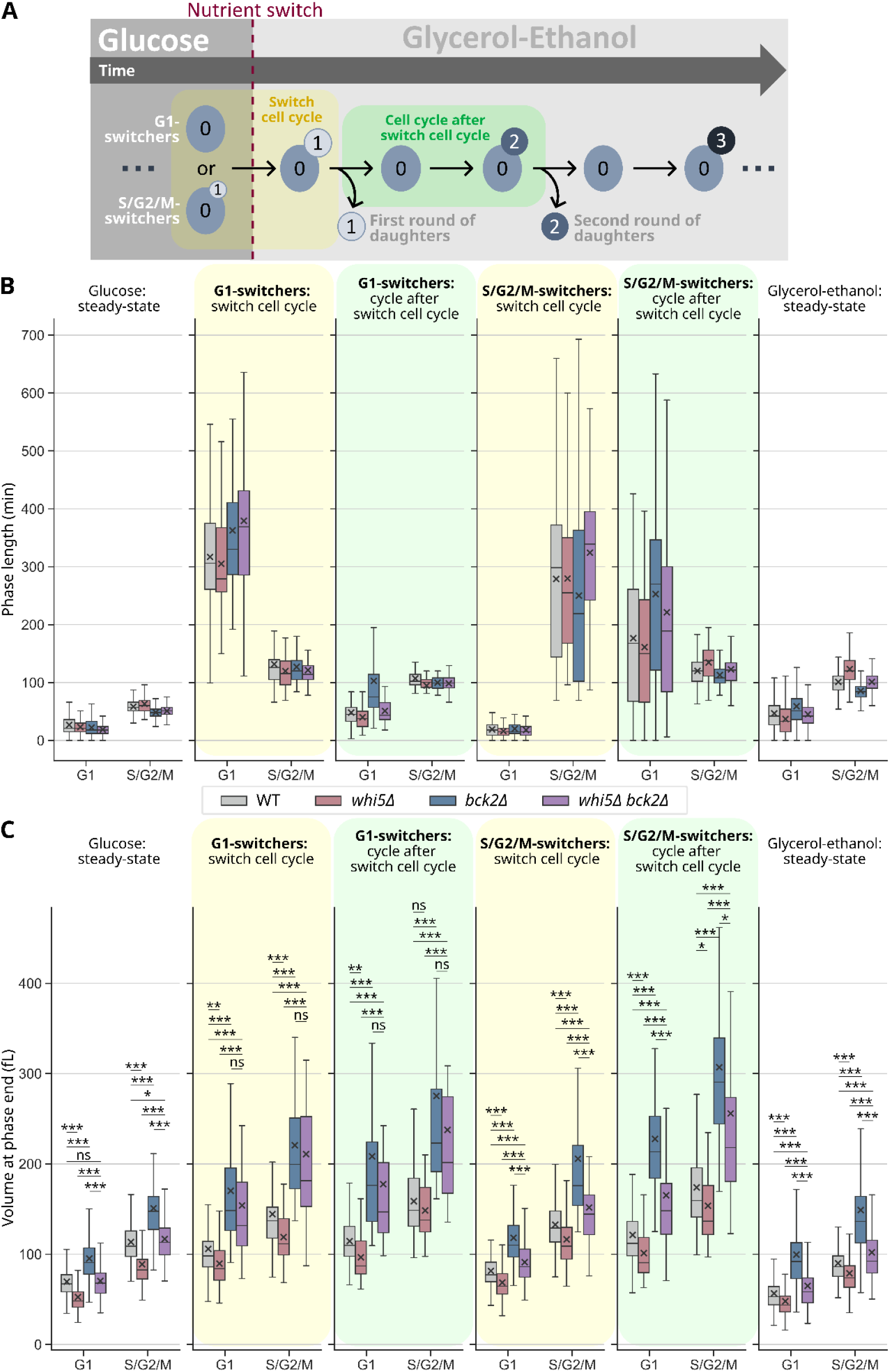
The nutrient switch causes cell cycle arrests in switchers and leads to stronger cell enlargement in *bck2Δ* cells. (**A**) Schematic explanation for cell categories analysed in this figure. (**B**) Phase lengths are plotted for *switchers’ switch cell cycles* (yellow) and *cell cycles after switch-cell-cycle* (green). x symbol denotes the mean phase length of the population. The outermost left and right panels show steady-state phase lengths for SCD and SCGE, respectively. For each growth medium, complete cell cycles were pooled from two independent steady-state experiments. The number of cells in each box in the steady-state panels is between 450 and 950. For the nutrient switch panels, cells were pooled from three independent experiments. For *G1-switchers’ switch cell cycle* G1 (yellow, left), n_WT_ = 144, n_*whi5Δ*_ = 59, n_*bck2Δ*_ = 67, n_*whi5Δbck2Δ*_ = 50. For *G1-switchers’ switch cell cycle* S/G2/M (yellow, left), n_WT_ = 132, n_*whi5Δ*_ = 46, n_*bck2Δ*_ = 56, n_*whi5Δbck2Δ*_ = 42. For *S/G2/M-switchers’ switch cell cycle* G1 (yellow, right), n_WT_ = 239, n_*whi5Δ*_ = 107, n_*bck2Δ*_ = 116, n_*whi5Δbck2Δ*_ = 115. For *S/G2/M-switchers’ switch cell cycle* S/G2/M (yellow, right), n_WT_ = 197, n_*whi5Δ*_ = 66, n_*bck2Δ*_ = 66, n*whi5Δbck2Δ* = 94. For *G1-switchers’ cell cycle after switch-cell-cycle* G1 (green, left), nWT = 120, n*whi5Δ* = 34, n*bck2Δ* = 49, n_*whi5Δbck2Δ*_ = 34. For *G1-switchers’ cell cycle after switch-cell-cycle* S/G2/M (green, left), nWT = 91, n_*whi5Δ*_ = 14, n_*bck2Δ*_ = 39, n_*whi5Δbck2Δ*_ = 26. For *S/G2/M-switchers’ cell cycle after switch-cell-cycle* G1 (green, right), nWT = 164, n_*whi5Δ*_ = 47, n_*bck2Δ*_ = 51, n_*whi5Δbck2Δ*_ = 61. For *S/G2/M-switchers’ cell cycle after switch-cell-cycle* S/G2/M (green, right), n_WT_ = 138, n_*whi5Δ*_ = 45, n_*bck2Δ*_ = 29, n_*whi5Δbck2Δ*_ = 46. (**C**) This dataset is also used to determine cell volume at the end of the respective cell cycle phases (fL). Cell volume at the end of S/G2/M is a sum of mother and bud volume at the last frame before division. Independent two-tailed t-tests assuming unequal variances (Welch’s t-tests) were used for statistical analyses.

For each strain, the immediate G1 arrest of *G1-switchers* led to an increased cell volume at the end of G1 (Fig. 4C, left yellow). The following G1, although short, slightly increased the cell volume further (Fig. 4C, left green). Similarly, also for *S/G2/M-switchers* of all strains, the G1 arrest in the *cell-cycle-after-switch-cell-cycle* led to cell enlargement (Fig. 4C, right green). During these G1 arrests, strains that lacked Bck2 exhibited stronger enlargement than wild-type and *whi5Δ* cells in both categories of *switchers* (Fig. 4C). These G1 arrests, therefore, were sufficient to cause significant differences in cell volume at the end of G1 between wild-type and *whi5Δbck2Δ* cells. The G1 arrests also contribute to the diverging CV and mean cell volume observed post-switch in the bulk nutrient switch experiment (Fig. 2).

### Cells that face the nutrient switch as buds arrest in their first G1, leading to stronger cell enlargement in *bck2Δ* cells

So far, we have shown that deletion of *BCK2* leads to more enlarged *switchers* during their first G1 after the switch. Apart from the actual *switchers*, also the *first round of daughters* of *S/G2/M-switchers* faced the switch because they were present as buds (Fig. 5A, highlighted in blue, cell number 1). The *first round of daughters of G1-switchers*, on the other hand, first appeared after the nutrient switch. G1 duration was 3-to 4-fold longer for the *first round of daughters* of *S/G2/M-switchers*, indicating that they arrested in G1 (Fig. 5B). This phenomenon was specific for this category of cells, as daughter cells born in the following rounds and those of *G1-switchers* showed G1 durations more similar to steady-state growth on SCGE (Fig. 5B). Consequently, the *first round of daughters of S/G2/M switchers* was also larger at G1 end, with *bck2Δ* and *whi5Δbck2Δ* cells again showing significantly stronger enlargement than the other strains (Fig. 5C). The G1 arrest was also longer for the Bck2 mutant strains as compared to the others (Fig. 5B). It appears that the cells that faced the switch in S/G2/M, either as mother or bud, carried the memory of the switch into the next cell cycle, leading to a prolonged G1 phase in the next cycle. Moreover, the first daughters of S/G2/M switchers showed weakened size control (Fig. S4) and lower survival (Fig. S5).

**Figure 5.**
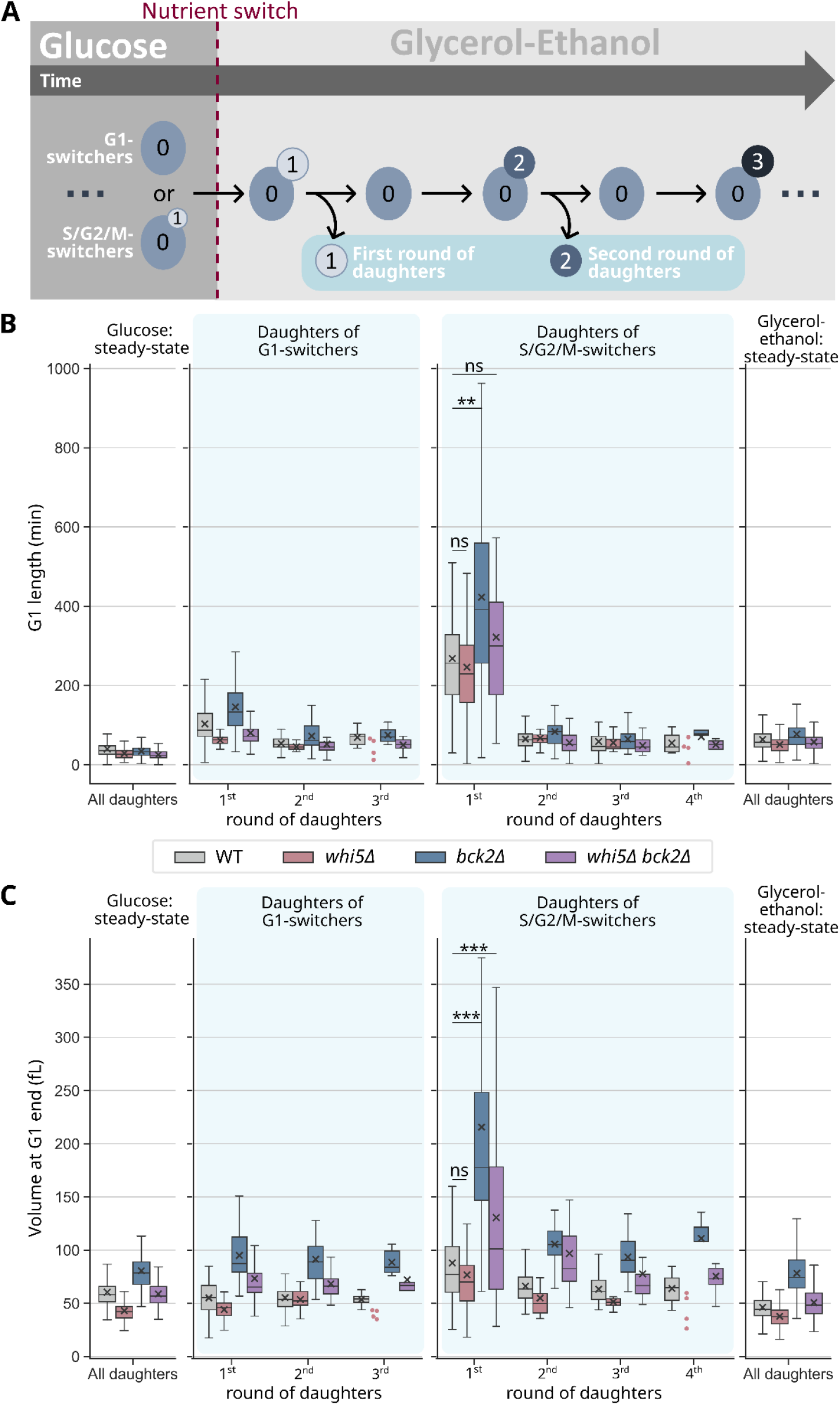
Cells that face the nutrient switch as buds arrest in their first G1, leading to stronger enlargement in *bck2Δ* cells. (**A**) A schematic explanation for cell populations depicted in this figure. (**B**) G1 lengths are plotted for the *rounds of daughters* (blue) of *G1-switchers* and *S/G2/M-switchers*. x symbol denotes the mean phase length of the population. The outermost left and right panels show steady-state phase lengths for SCD and SCGE, respectively. For each growth medium, complete cell cycles of daughter cells were pooled from two independent steady-state experiments. The number of cells in each box in the steady-state panels is between 198 and 330. For the nutrient switch panels (blue), cells were pooled from three independent experiments. For the *first round of daughters of G1-switchers*, n_WT_ = 103, n_*whi5Δ*_ = 21, n_*bck2Δ*_ = 48, n_*whi5Δbck2Δ*_ = 33. For the *first round of daughters of S/G2/M-switchers*, n_WT_ = 123, n_*whi5Δ*_ = 40, n_*bck2Δ*_ = 34, n_*whi5Δbck2Δ*_ = 50. The next *rounds of daughters* had fewer cells than the first *round*. If a category includes less than 5 cells, the individual data points are shown. (**C**) The same dataset was also used to determine cell volume at the end of the G1 (fL). Independent two-tailed t-tests assuming unequal variances (Welch’s t-tests) were used for statistical analyses.

### *First round of daughters* of *S/G2/M-switchers* arrests mainly in pre-*Start* G1

After showing that *S/G2/M-switchers* and their first daughters exhibit a prolonged first G1 following the nutrient switch, we asked at what point in G1 cells arrest, and whether this arrest depends on the exact cell cycle stage at the time of the nutrient switch. To further characterise this multi-generational arrest-phenotype, we therefore employed cell cycle reporters that allowed us to resolve the cell cycle into more specific cell cycle phases. In particular, we used an mCitrine-tagged mutant *WHI5* allele, *WHI5-WIQ*, integrated into the *URA3* locus while the endogenous *WHI5* gene remained unaltered. The Whi5-WIQ protein is a loss-of-function mutant that does not bind SBF ^40^ but retains its ability to localise to the nucleus in a cell-cycle-dependent manner. In addition, we tagged the histone *HTB2* with the fluorescent protein mScarlet-I. The deletions of interest *–* Δ*whi5*, Δ*bck2*, and Δ*whi5*Δ*bck2 –* were then introduced into this cell cycle-reporter strain and the nutrient switch experiment was repeated. Fluorescently tagged Htb2 allowed us to segment a nuclear mask for each cell. Using the nuclear mask along with cellular segmentation, we could quantify fluorescence signal in three compartments: the whole cell, the nucleus, and the cytoplasm. Whi5 has previously been reported to be nuclear from telophase until *Start*, while CDK activity is low ^41^. Fig. 6A shows example images, as well as quantified Whi5-WIQ-mCitrine and HTB2-mScarlet-I signal traces for one steady-state (SCD) example cell in yellow and red, respectively. For cell categories of interest, we identified *Start* as the G1 frame at which 50% of Whi5 had exited the nucleus ^42^(Fig. 6A). We used the Htb2-mScarlet-I amount in the bud as a marker for the entry of the dividing nucleus into the bud, which occurs during anaphase in mitosis ^43^(Fig. 6A). Anaphase was presumed to continue until Whi5-WIQ-mCitrine re-entered the nucleus ^41^(Fig. 6A). The nuclear re-entry of Whi5-WIQ was annotated as the start of telophase and subsequent frames were labelled as post-anaphasic (Fig. 6A). Fig. 6B shows the cell cycle phases of the *first round of daughters* of *S/G2/M-switchers*. After splitting G1 into pre- and post-*Start*, we found that the G1 arrest observed in the *first round of daughters* of *S/G2/M-switchers* was predominantly in pre-S*tart* G1 (dark green areas, Fig. 6B). Upon quantification (Fig. 6C), we found that *bck2Δ* and *whi5Δbck2Δ* cells had longer pre-*Start* G1 durations as compared to wild type and *whi5Δ*.

**Figure 6.**
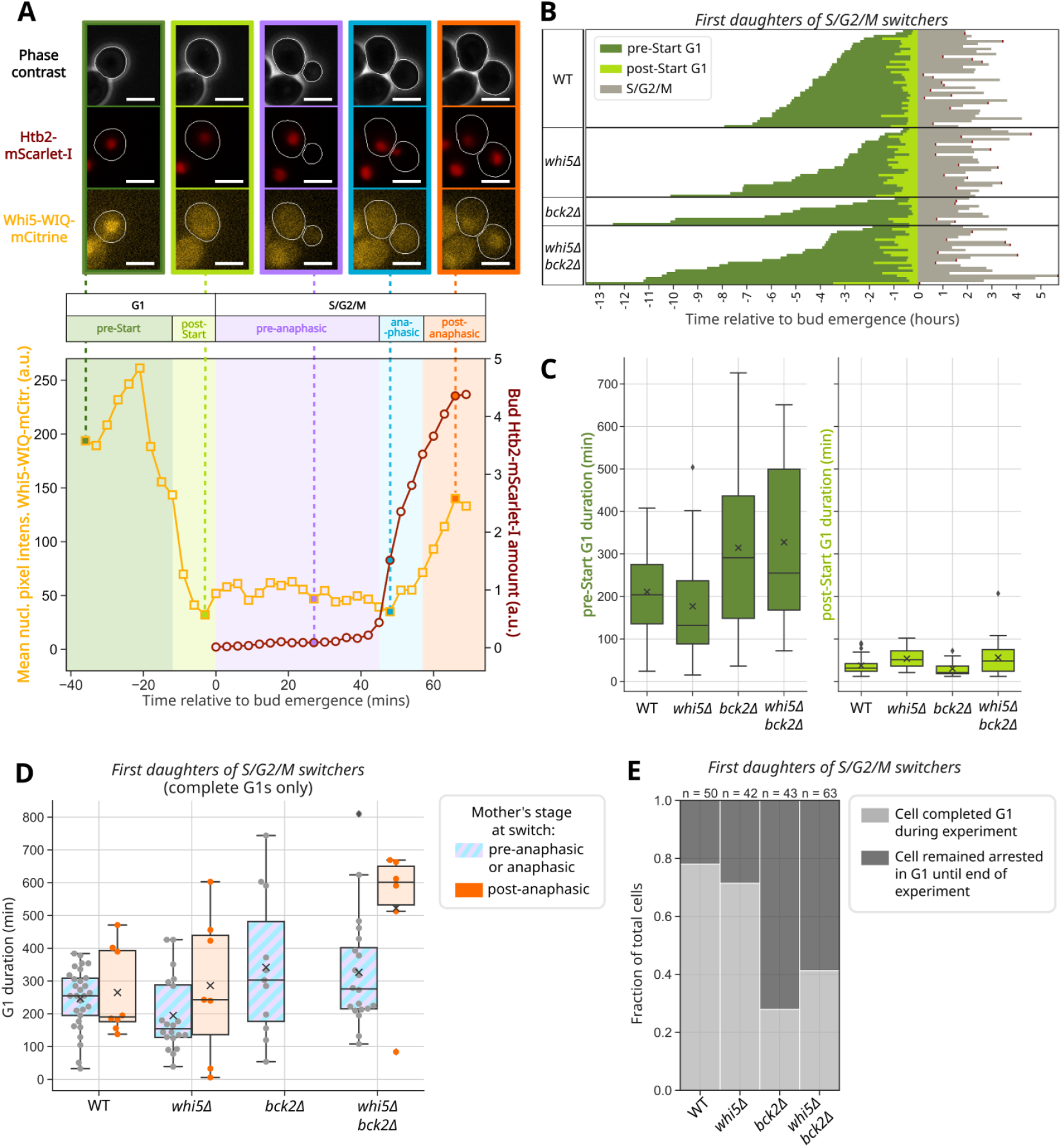
*First round of daughters* of *S/G2/M-switchers* arrest mainly in pre-Start G1. In *bck2Δ* mutants, the strongest arrest phenotypes are observed in first daughters whose mothers faced the nutrient switch after anaphase. (**A**) An example of cell cycle phase resolution for a steady-state (SCD) wild-type cell based on two fluorescent signals: mean nuclear pixel intensity of Whi5-WIQ-mCitrine (yellow trace, a.u.) and bud Htb2-mScarlet-I amount (red trace, a.u.), both plotted against time relative to bud emergence. 50% Whi5-WIQ-mCitrine nuclear exit was annotated as *Start*. Entry of Htb2-mScarlet-I into the bud was annotated as start of anaphase. Re-entry of Whi5-WIQ-mCitrine into the nucleus was annotated as end of anaphase. Dotted lines connect example images to the time-point at which they were acquired. Example images show cells in phase contrast (grey), histone Htb2 tagged with mScarlet-I (red) and Whi5-WIQ tagged with mCitrine (yellow). Cell contours generated with YeaZ in the Cell-ACDC pipeline are shown in white. Scale bars represent 4 µm. (**B**) Cell cycles of *first daughters of S/G2/M-switchers* with complete G1 phases were resolved into pre-Start G1 (dark green), post-Start G1 (light green) and S/G2/M (grey). A categorical heatmap shows cell cycle phases of single cells (as rows), plotted against time relative to bud emergence. Incomplete S/G2/M phases are capped with red ends. Black horizontal lines separate cells of different strains. (**C**) A quantification of pre-Start G1 durations and post-Start durations is plotted as a box-plot. x symbol denotes the mean phase duration of the population. Data was pooled from two independent experiments of live-cell microscopy coupled to nutrient switch. n_WT_ = 38, n_*whi5Δ*_ = 27, n_*bck2Δ*_ = 11, n_*whi5Δbck2Δ*_ = 23. The durations of complete G1 phases of *first daughters of S/G2/M-switchers* are plotted in (**D**). The *first daughters of S/G2/M-switchers* are further categorised on the basis of their mothers’ cell cycle phase at the time of nutrient switch: pre-anaphasic or anaphasic (purple and blue striped) or post-anaphasic (orange). x symbol denotes the mean phase duration of the population. (**E**) The fraction of first daughters of S/G2/M-switchers that complete G1 during the experiment (light-grey) and those that stay arrested in G1 until the end of the experiment (cells with incomplete G1 phases, dark-grey) are plotted. Incomplete G1 phases are G1 phases interrupted by the end of the experiment and not by death or exclusion of cells from the analysis.

### In *bck2Δ* cells, the first daughters of mothers which faced the nutrient switch after anaphase have the strongest arrest phenotypes

Next, we asked whether the specific cell cycle phase of *S/G2/M-switchers* at the time of the nutrient switch affects the severity of the G1 arrest in their *first round of daughters*. Thus, we categorised the *first round of daughters* of *S/G2/M-switchers* by whether the mother was pre-anaphasic, anaphasic or post-anaphasic at the time of nutrient switch, and analysed the corresponding G1-lengths (Fig. 6D, Fig. S6). Fig. 6D shows the durations of the G1 phases of the first round of daughters of *S/G2/M-switchers* that completed G1 during the course of the experiment. The fraction of the first round of daughters of *S/G2/M-switchers* that did not complete G1, *i*.*e*., remained arrested in G1 until the end of the experiment, is shown in Fig. 6E and the lower limits of their G1 durations are plotted in Fig. S6. For *whi5Δbck2Δ* cells, the strongest arrest phenotype, *i*.*e*., the longest G1 lengths, were observed for the group whose mothers were post-anaphasic at the time of nutrient switch (Fig. 6D). For *bck2Δ*, we did not observe any complete G1 phases for the group of daughters whose mothers were post-anaphasic at the time of nutrient switch (Fig. 6D) as all of these daughters stayed arrested in G1 until the end of the experiment (Fig. S6, Fig. 6E). In fact, for *bck2Δ* and *whi5Δbck2Δ* cells, the majority of the first round of daughters of *S/G2/M-switchers* did not complete G1 until the end of the experiment (Fig. 6E). Thus, our data suggest that Bck2 has a nutrient switch-specific function after anaphase which affects G1 exit in the next generation. Such a role for Bck2 had previously been proposed ^21^, where Bck2 was speculated to integrate environmental information into the cell cycle progression decision at multiple cell cycle-phase transitions, including M/G1.

Additionally, it has previously been shown that Bck2 promotes cell cycle progression through *Start* by promoting the expression of Cln3, Swi4 and Cln2 ^21^. It was also shown that Cln3 is a short-lived protein and depleted after a switch from a rich to a poor carbon source ^26^ and that *cln3Δbck2Δ* is inviable due to permanent G1 arrest. One explanation for the extended G1 arrests in Bck2 deletion mutants we identified here could therefore be a post-switch Cln3 depletion, which, coupled with the absence of Bck2, temporarily mimics a *cln3Δbck2Δ* phenotype.

### Bck2, but not Cln3, is critical for size adaptation after a nutrient switch

This led us to ask if similar to *bck2Δ, cln3Δ* also has a strong arrest phenotype after a nutrient switch, or if, as would be expected if Cln3 is depleted, Cln3 is dispensable for cell size adaptation after the switch. During steady-state growth on SCD, *cln3Δ* cells are larger than wild-type cells, similar to *bck2Δ* (Fig. 7A). The large size of *bck2Δ* and *cln3Δ* can be partially rescued by an additional deletion of Whi5 (Fig. 7A). The *whi5Δbck2Δcln3Δ* triple deletion is similar in size to the *bck2Δ* and *cln3Δ* single deletions (Fig. 7A), which further highlights that the effects of the Whi5, Bck2 and Cln3 deletions are additive.

**Figure 7.**
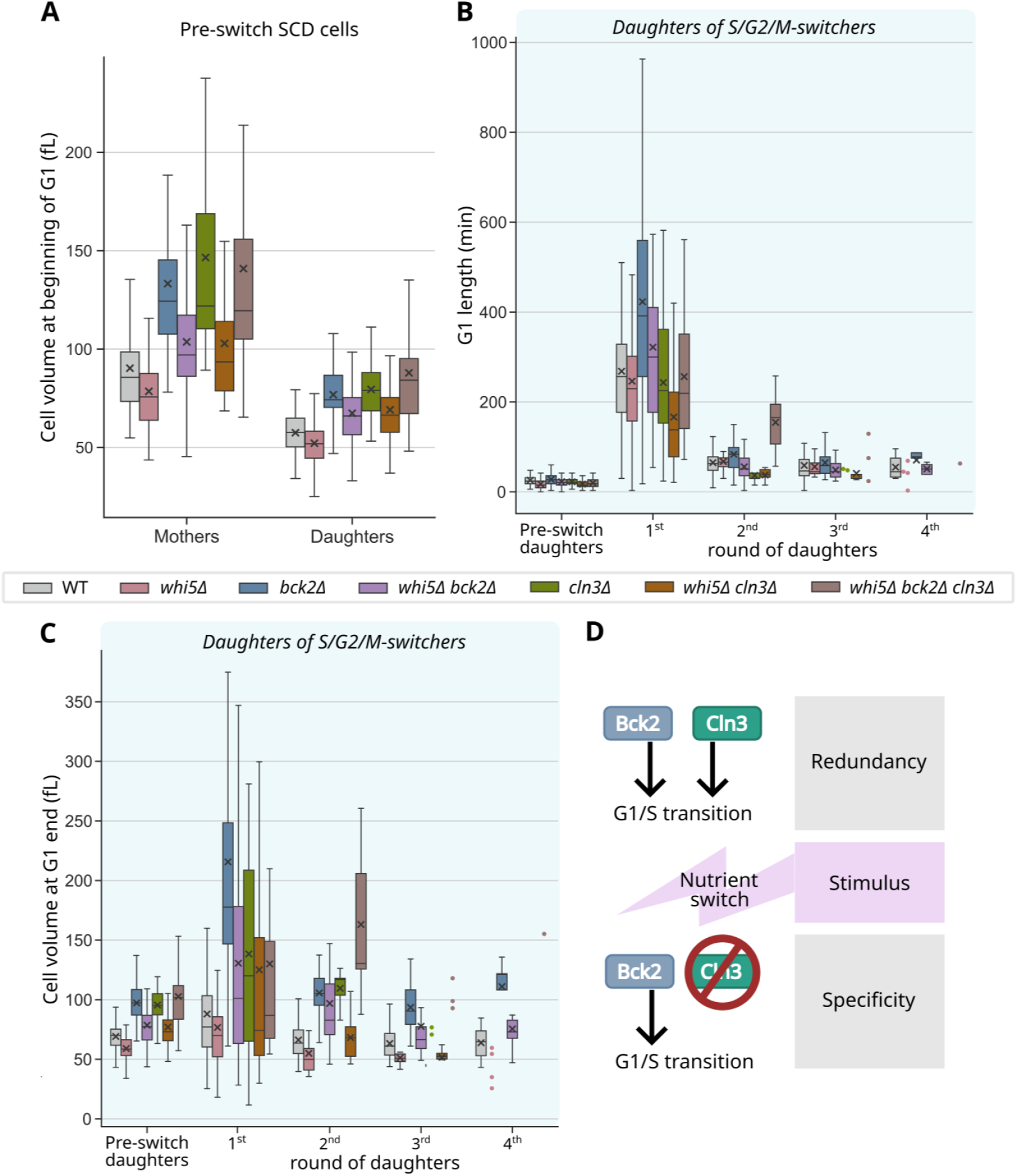
*bck2Δ* cells have a longer G1 arrest and stronger enlargement than *cln3Δ* cells after a nutrient switch. (**A**) Cell volume at the beginning of G1 is plotted for daughter (first-generation) and mother (generation>1^st^) cells of various strains. x symbol denotes the mean phase duration. For daughters, n_WT_ = 164, n_*whi5Δ*_ = 82, n_*bck2Δ*_ = 73, n_*whi5Δbck2Δ*_ = 77, n_*cln3Δ*_ = 44, n_*whi5Δcln3Δ*_ = 55, n_*whi5Δbck2Δcln3Δ*_ = 37. For mothers, n_WT_ = 209, n_*whi5Δ*_ = 87, n_*bck2Δ*_ = 95, n_*whi5Δbck2Δ*_ = 95, n_*cln3Δ*_ = 48, n_*whi5Δcln3Δ*_ = 61, n_*whi5Δbck2Δcln3Δ*_ = 34. (**B**) G1 lengths and (**C**) cell volumes at G1 end are plotted for the *rounds of daughters* of *S/G2/M-switchers*. The x symbol denotes the mean. Pre-switch daughters are daughter cells born during steady-state in SCD before the switch. The dataset for wild type, *whi5Δ, bck2Δ*, and *whi5Δbck2Δ* is the same as in Figure 5. For *cln3Δ, whi5Δcln3Δ* and *whi5Δ bck2Δcln3Δ*, cells were pooled from two independent experiments. For the *first round of daughters of S/G2/M-switchers*, n*cln3Δ* = 16, n*whi5Δcln3Δ* = 39, n*whi5Δbck2Δcln3Δ* = 11. The next *rounds of daughters* had fewer cells than the first *round*. If a category includes less than 5 cells, individual data points are shown.

In contrast, during adaptation to the nutrient switch, *cln3Δ* cells do not have as strong a phenotype as *bck2Δ* cells, both in terms of the G1 length of the first *daughters of S/G2/M switchers* (Fig. 7B) and their volume at G1 end (Fig. 7C). While in steady-state, the *whi5Δbck2Δcln3Δ* triple deletion resembles the *bck2Δ* and *cln3Δ* single deletions in size (Fig. 7A), in the first daughters of S/G2/M switchers born after the nutrient switch, its size is closer to the *whi5Δbck2Δ* double deletion, not the *bck2Δ* single deletion (Fig. 7C). This observation is consistent with a nutrient switch associated depletion of Cln3, rendering Bck2 critical during the nutrient switch. Moreover, it highlights that unique functions for seemingly redundant proteins can be uncovered by studying cells in dynamically changing environments.

## Discussion

In nature, yeast cells regularly face changing environments. Cell size control therefore comprises two tasks: size homeostasis, which is the maintenance of narrow size distributions in largely constant environments, and size adaptation, which is the adjustment of cell size to the optimum for a changed environment. So far, most studies on cell size regulation have focused on steady-state conditions. While these studies have been useful for identifying size homeostasis mechanisms and size regulators, they cannot, by design, provide insights into mechanisms of size adaptation. This suggests that studying cell size adaptation in response to nutrient switches may reveal aspects of cell size regulation that were not accessible through steady-state experiments. In particular, we asked whether nutrient switches could give insights into specific functions of cell size regulators that appear redundant in steady-state size control.

The analysis of nutrient switch experiments at the single-cell level is even more complex than that of steady-state experiments. In steady-state, key insights can be gained by analysing individual cell cycles, without tracking cells over multiple generations. In nutrient switch experiments, the history of a cell, specifically the cell cycle stage at the time of the nutrient shift, affects how the cell and its offspring adapt to the new environment (this study and ^19^). To understand cell size adaptation, it is, therefore, crucial to categorise cells based on their cell cycle stage at the time of the nutrient switch before following their progeny over multiple generations. Several studies have investigated nutrient switches with live-cell microscopy in different contexts ^19,44–47^. However, none of them performed multi-generational lineage tracking in addition to cell categorisation based on the cell cycle stage at the time of the nutrient switch. Here, we built on recent progress in machine learning approaches and performed the required analysis using the image analysis tools YeaZ and Cell-ACDC ^38,39^. This enabled us to track complete cell lineages over multiple generations throughout the adaptation to a nutrient switch and create a framework for dealing with the complexity of single-cell categorisation.

From bulk experiments, we found that while the simultaneous deletion of two important size regulators, Whi5 and Bck2, hardly affects cell size or cell-size homeostasis during steady-state growth in glucose media, it disrupts both after a nutrient switch from glucose to glycerol/ethanol media. Around 10 to 25 hours following a nutrient switch, both wild-type and *whi5Δbck2Δ* cells show a strong increase in the CV of cell volume, indicating weaker size homeostasis. This disruption of size-homeostasis is stronger in the *whi5Δbck2Δ* strain. Consistent with our findings, a similar post-switch increase in the CV of cell volume has also been observed in a recent study that analyses budding yeast adaptation to a nitrogen-downshift ^46^. Using the cell history-based categorisation explained above, we followed single cells through the nutrient switch and found that *bck2Δ* mutants show longer cell cycle arrests after the nutrient switch. Cells that were already budding at the time of the nutrient switch as well as their first daughters born after the nutrient switch exhibit the longest cell cycle arrests, the strongest cellular enlargement and the lowest survival post-switch. The following rounds of daughters closely resemble those in exponential growth in terms of cell cycle phase durations and cell size. In contrast to steady-state, where deletion of the G1/S activators *BCK2* and *CLN3* lead to comparable size phenotypes, *cln3Δ* cells show a much weaker phenotype in response to the nutrient shift. Moreover, an additional deletion of *CLN3* in the *whi5Δbck2Δ* strain does not affect the post-switch arrest or enlargement phenotype. Taken together, our findings suggest that while Cln3 and Bck2 have an apparently redundant role in steady-state size control, Bck2 fulfils a specific function during nutrient adaptation. One intriguing explanation for this is that Cln3 may be depleted after the nutrient switch, as observed by Sommer *et al*. ^26^. We therefore propose that the shared role for Bck2 and Cln3 in activating the G1/S transition serves as a potential ‘backup plan’ for G1-exit when Cln3 is depleted due to changing nutrient environments (Fig. 7D).

It is also known that Bck2 interacts with multiple proteins involved in nutrient sensing and promotes expression of cell cycle regulators ^21^. One of the proteins it interacts with is Tpd3, which is the scaffolding subunit of the protein phosphatase PP2A ^21^. PP2A is part of the cascade by which two nutrient-sensing pathways, TOR and PKA, regulate *Start* ^48–50^. This interaction places Bck2 directly at the link between nutrient-dependent growth signalling and cell cycle progression. Thus, a role of Bck2 in nutrient signalling could also contribute to the longer cell cycle arrests and stronger cellular enlargement observed in *bck2Δ* cells after a nutrient switch event.

In conclusion, our work here provides new insights into size adaptation to nutrient challenges. We show that the redundancy between the roles of Bck2 and Cln3 at the G1/S transition is specific to steady-state growth and Bck2 becomes the more crucial G1/S transition activator following a nutrient switch (Fig. 7D). More generally, our work suggests that studying cell size regulation in conditions other than steady-state is key to revealing unique roles for apparently redundant size regulators. We, therefore, expect that using similar approaches for a broader range of mutants and perturbations in yeast as well as other organisms will give new mechanistic insights into the processes underlying size homeostasis and adaptation.

## Methods

### Yeast strains

All *Saccharomyces cerevisiae* strains used in this study are haploid derivatives of W303. They were constructed using standard methods and verified by sequencing. A full genotypic description of the strains is available in Supplementary table 1. Yeast strains are available upon reasonable request.

### RNA extraction and RT-qPCR

Cells were grown in 50 ml SCD for at least 17 hours and maintained at OD_600nm_ <1 before harvest. At OD_600nm_ > 0.1, total RNA was extracted using the RNA extraction protocol of the YeaStar RNA Kit (Zymo Research). The RNA was treated with Turbo DNAase enzyme (Thermo Fisher Scientific) as per the manufacturer’s protocol. The quality of the RNA was examined in an RNA gel and the concentration and purity were checked using the NanoDrop OneC spectrophotometer from Thermo Fisher Scientific. 1 µg of RNA was reverse transcribed with random primers and the high-capacity cDNA reverse transcription kit from Thermo Fisher Scientific. The resulting cDNA was diluted 1:10 fold using double-distilled water. Quantitative PCR (qPCR) was performed using 2 µl of these dilutions as template. Target-specific primers and SsoAdvanced Universal SYBR Green Supermix (BioRad) were used for qPCR reactions in 96-well plates (Roche LightCycler 480 Multiwell Plate 96). For each biological replicate, mean Cq values of target genes were calculated by averaging across at least three technical replicates. The mean Cq value of the reference gene, *ACT1*, was subtracted from the mean Cq value of *WHI5* to calculate normalised *WHI5* Cq values (ΔCq). The *WHI5* ΔCq value of wild type was then subtracted from that of the strains of interest to obtain the *WHI5* ΔΔCq value for each strain. The relative *WHI5* mRNA concentration was calculated using the following formula

xs

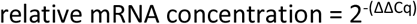

This process was repeated for five different RNA extractions (experimental replicates). These were plotted as a boxplot in Fig. 1B.

### Single molecule fluorescence in situ hybridisation (smFISH)

smFISH was performed using WHI5-mRNA specific Stellaris® FISH probes from Biosearch Technologies. The targeted probes consisted of 37 18 nucleotide long oligonucleotides, each labelled with the Quasar-570® dye. The smFISH protocol for *S. cerevisiae* described on www.biosearchtech.com/stellarisprotocols was used for sample preparation. Cells were grown for a minimum of 17 hours in 50 ml SCD and maintained at OD_600nm_ <1. At an OD_600nm_ between 0.3 and 0.5, 45 ml of these cultures were fixed by application of 4% formaldehyde (final concentration) for a period of 45 minutes. The cells were centrifuged at 1600 g for 4 minutes and washed twice, each time with 1 ml of ice-cold fixation buffer comprising 1.2 M sorbitol (SigmaAldrich) and 0.1 M K_2_HPO_4_ (Sigma-Aldrich) at pH 7.5. Cells were then incubated in 1 ml of fixation buffer containing 6.25 µg Zymolyase (Biomol) for 55 minutes at 30 °C. After two more washes with ice-cold fixation buffer, the fixed and digested cells were stored overnight in 70% ethanol at 4 °C. On the next day, 300 µl of cells in 70% ethanol were centrifuged (400 g, 5 minutes) and resuspended in 100 µl of Stellaris® RNA FISH hybridization buffer (Biosearch Technologies) containing 10% v/v formamide and 125 mM smFISH probes. These cells were incubated overnight in the dark at 30 °C to allow for hybridisation of smFISH probes to mRNA. After overnight hybridisation, cells were washed with Stellaris® RNA FISH wash buffer A (Biosearch Technologies) containing 10% v/v formamide and resuspended in 1 ml Stellaris® RNA FISH wash buffer A (Biosearch Technologies) containing 10% v/v formamide and 5 ng/ml DAPI stain. Cells were incubated in the DAPI staining solution for 30 minutes at 30 °C, before being washed with Stellaris® RNA FISH wash buffer B (Biosearch Technologies). Washed cells were mounted onto glass slides in Vectashield® mounting medium (Vector Laboratories) for the first replicate and in ProLong Gold mounting medium (Thermo Fisher Scientific) for the second replicate due to unavailability of Vectashield®. The prepped smFISH samples were imaged on a Zeiss LSM 800 microscope using a 63×/1.4 NA oil immersion objective and an Axiocam 506 camera. The Zen 2.3 software was used to acquire multicolor z-stacks containing 20-25 z-slices at 240 nm intervals. Bright field images were acquired with the TL LED using an exposure time of 140 ms at 6% of maximum intensity. For imaging Quasar-570®, an exposure time of 5 s to illumination by a 530 nm LED at 50% intensity was used. DAPI was imaged with an exposure time of 130 ms and illumination at 385 nm and 30% intensity.

### Data analysis for smFISH

Cells were segmented in bright field using the YeaZ neural network ^39^ in the Cell-ACDC interface ^38^. After manual correction of segmentation masks and annotation of bud-mother relationships in Cell-ACDC, the smFISH fluorescence spots were counted using a custom spot detection routine in python, described in detail in ^51^ and ^37^. For each strain and experimental replicate, the following settings of the custom routine were adjusted to optimise spot detection. The 3D Gaussian filter was applied with a sigma in the range of 0.3 to 2 voxels. The ‘threshold_triangle’ automatic thresholding algorithm was used for instance segmentation of the spot signal. The effsize_glass_s filter was used to filter for valid peaks using an effect size threshold in the range of 1 to 2.1. Spots were also filtered for size with the lower limit ranging from 1.6 to 2 pixels and the upper limit ranging from 4 to 20 pixels.

Spot detection was followed by categorisation of cells into one of three different cell cycle-stages: G1, S and G2/M. Unbudded cells with a single nucleus were categorised into G1 stage. Budded cells with a bud volume-to-mother volume ratio of <0.3 were categorised as S stage. The remaining budded cells were categorised as G2/M cells. Since Whi5 transcription peaks in S-phase, Fig. 1C shows the average number of Whi5 mRNA in S-phase cells.

### Bulk nutrient switch experiments

Yeast cells were inoculated into 3 mL Yeast Peptone Medium containing 2% Glucose (YPD medium) and incubated at 30 °C in a shaking incubator at 250 rpm (Infors, Ecotron) for at least 4 hours. After four hours, cells were washed and inoculated into 50 mL synthetic complete medium with 2% glucose (SCD medium) at a starting OD_600nm_ of 0.0001 and incubated overnight at 30 °C under shaking conditions (250 rpm). Cells were grown for a minimum of 17 hours in 50 ml SCD and maintained at OD_600nm_ <1. At OD_600nm_ between 0.5 and 0.6, cells were washed with and inoculated in synthetic complete medium with 2% glycerol and 1% ethanol (SCGE medium) at a starting OD _600nm_ of 0.01 and incubated under the same conditions (30 °C, 250 rpm). Starting from the time of inoculation in SCGE medium, OD_600nm_ and coulter-counter measurements were performed at 1-1.5 hour intervals until 28.5 hours had passed since the nutrient switch. The same measurements were also performed after 48 hours and 72 hours had passed since the nutrient switch. For the measurements taken before the 30-hour time point, culture OD_600nm_ was always below 1. For measurements taken 48 and 72 hours after the nutrient switch, once the cultures grew to an OD_600nm_ of 0.1, appropriate dilutions were made to maintain the OD_600nm_ between 0.1 and 1. The complete time course of measurements was achieved in sections in three different experiments of two replicates each. Therefore, overlapping time points in the plot have four data points. The CV (coefficient of variation) was calculated from the coulter-counter size distribution with the formula:

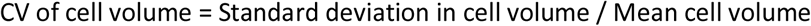

For statistically comparing the two strains, we divided the time course into three ten-hour intervals and grouped the remaining time points together (Fig. 2B, D). For each time-interval, we performed a mixed ANOVA test, with time as the within-subjects factor and strain as the between-subjects factor for classification. The interaction between the two factors was not significant, and therefore we could interpret the effects of each factor individually. The p-values obtained for each time interval based on the between-subjects factor *i*.*e*., strain, were represented in statistical annotations in the figure (Fig. 2B, D).

### Live-cell microscopy

#### Culture conditions

Yeast cells were inoculated into 3 mL Yeast Peptone Medium containing 2% Glucose (YPD medium) and incubated at 30 °C in a shaking incubator at 250 rpm (Infors, Ecotron) for at least 5 hours. After 5 hours, cells were washed and inoculated into 50 mL synthetic complete medium with either 2% glucose (SCD medium) or 2% glycerol and 1% ethanol (SCGE medium). The cells were allowed to grow for at least 16 hours and maintained at OD_600nm_ <1. They were then diluted to OD_600nm_ = 0.1 for microscopy.

#### Microscopy setup

Live-cell time-lapse microscopy was performed using a Nikon Eclipse Ti-E microscope in combination with a custom microfluidic device, described in detail in ^37^. The microfluidic device consisted of a polydimethylsiloxane replica with functional structures bonded to a glass cover slip. This created a cavity that could trap cells and limit colony growth to the *XY*-plane. Growth medium was pumped through the microfluidic device at a constant rate of 20 µL/min to sustain the growth of cells in the device during the imaging process. For steady-state experiments, filtered SCD or SCGE medium was used for the entire duration of the experiment. For nutrient switch experiments, cells were grown in SCD for the first two hours of the experiment and then the medium was switched to SCGE for the rest of the experiment. The time taken for the medium to be completely switched within the device was calculated to be 30 minutes and medium switch was therefore annotated at 2.5 hours in the downstream analysis of the live-cell microscopy data. To maximise the rounds of daughters observed per video before crowding of the field of view prevented further analysis, the microscopy experiments were started with only one or two cells per position. This limited the number of cells that could be analyzed at the time of the nutrient switch, which were further reduced by repeated categorisation.

The epifluorescence microscopy set-up included the Nikon Eclipse Ti-E microscope with NIS-Elements software, SPECTRA X light engine illumination and the Andor iXon Ultra 888 camera. Phase-contrast and fluorescence images were acquired at 3-minute intervals via a plan-apo λ 100×/1.45 Na Ph3 oil immersion objective. An additional magnification of 1.5x was applied at the time of imaging. The phase contrast images were acquired using an exposure time of 100 ms. For mCitrine fluorescence imaging, an exposure time of 300 ms was used and the SPECTRA X light engine was used for illumination at 508 nm and 40% power (24.8 mW). mScarlet-I fluorescence was imaged with an exposure time of 200 ms and illumination at 555 nm and 10% power (26 mW). The image acquisition settings were the same across all live-cell microscopy experiments. The incubation temperature of the cells during live-cell microscopy was maintained at 30 °C using the objective heater and a custom heatable insertion.

### Data analysis for live-cell microscopy

Live-cell microscopy data was analysed with the open-source Cell-ACDC software ^38^. In the data preparation step, the images were aligned and cropped to the region of interest. Automated segmentation and tracking was performed on the phase contrast channel using the YeaZ neural network ^39^ in the Cell-ACDC interface. The resulting segmentation masks were manually corrected and pedigrees were assigned in the Cell-ACDC GUI. The annotated and manually corrected dataset could then be exported for downstream analysis.

#### Calculation of mean cell volume and CV of cell volume

For the mean cell volume and CV of cell volume plots (Fig. 1F and 1G respectively), live cells were pooled from the last frames of all imaging positions per strain, to maximise number of cells in the analysis. The mean cell volume and its CV were calculated from this pooled population. In Fig. 1G, the bar plot heights are the CV values obtained from the pooled data and the •, ▀, and **x** symbols show CV values obtained if the pooled data is split into the two underlying experimental replicates. To statistically compare the CV values between strains, bootstrapping was used to randomly sample positions from each strain in 10,000 iterations with replacement. For each combination of positions, live cells in the last frame were pooled and the CV of cell volume was calculated. To compare the bootstrapped CV distributions between strains, the vector of the CV distribution of one strain was subtracted from the vector of the CV distribution of the other. The 95%, 99% and 99.9% confidence intervals were calculated for the thereby obtained difference vector. If the confidence interval of the difference vector contained 0, it was concluded that the CV distributions overlapped considerably for the two strains at the given confidence level and their difference was not significant.

#### Calculation of cell cycle properties

Cell-cycle annotations in Cell-ACDC categorised unbudded cells into the “G1” phase and all budded cells into an umbrella category-the “S/G2/M” phase. The following functions from the Cell-ACDC downstream analysis Jupyter Notebook were used for calculating cell cycle properties such as phase lengths, and volume changes per phase and for retrieving relatives’ data for each cell: cca_functions.calculate_downstream_data, cca_functions.calculate_per_phase_quantities and cca_functions.calculate_relatives_data. From the resulting dataset, multi-generational mother-bud relationships, or pedigrees, could be acquired for cell populations of interest. Each cell cycle and cell cycle phase was labelled as complete or incomplete, based on whether the cell died during that cycle or phase and whether the cycle or phase was interrupted by the start or the end of the experiment. Unless specified otherwise, only complete cycles or complete phases were used for further analysis.

#### Assignment of cell cycle phases using cell cycle reporters

For the cell cycle-reporter strain, fluorescently labelled Htb2 allowed us to segment a nuclear mask for each cell. Using this along with cellular segmentation, we could quantify fluorescence signal in three compartments: the whole cell, the nucleus and the cytoplasm. Comparing Whi5-WIQ-mCitrine signal and Htb2-mScarlet-I signal between these compartments allowed us to further resolve the cell cycle into more specific phases than just G1 and S/G2/M, as described in the Results section. For budded cells, mean pixel intensity was calculated by summing up the sum pixel intensities of mother and bud as well as cell area in pixels for mother and bud and then dividing the total sum pixel intensity by the total area in pixels. The cytoplasm-adjusted nuclear mean pixel intensity of Whi5-WIQ-mCitrine (Fig. 6A) was calculated by subtracting cytoplasmic mean pixel intensity of Whi5-WIQ-mCitrine from nuclear mean pixel intensity of Whi5-WIQ-mCitrine. Htb2-mScarlet-I amount was background adjusted using a background ROI created during the data preparation step of the Cell-ACDC analysis pipeline (Fig. 6A).

For assigning the time point at which Start occurs, nuclear mean pixel intensity of Whi5-WIQ-mCitrine was plotted against time for each cell of interest. The frame at which nuclear mean pixel intensity of Whi5-WIQ-mCitrine started to decrease during G1 was manually identified (*Whi5_export_start_frame*). Similarly, the frame at which nuclear mean pixel intensity of Whi5-WIQ-mCitrine stopped decreasing was manually identified (*Whi5_export_end_frame*). The nuclear mean pixel intensities of Whi5-WIQ-mCitrine at these two frames were retrieved (*Whi5_intensity_export_start, Whi5_intensity_export_end*) and the difference between the two intensities was calculated. The difference was divided by 2 and added to *Whi5_intensity_export_end* to get nuclear mean pixel intensity of Whi5-WIQ-mCitrine when half of Whi5 has been exported from the nucleus (*Whi5_intensity_50%_export)*. Starting from *Whi5_export_start_frame* and proceeding frame-by-frame, the nuclear mean pixel intensity of Whi5-WIQ-mCitrine was compared to *Whi5_intensity_50%_export*. The first frame where frame-specific nuclear mean pixel intensity of Whi5-WIQ-mCitrine was less than *Whi5_intensity_50%_export* was assigned as the first post-Start frame. All following G1 frames were categorized as ***post-Start G1*** and all previous G1 frames were categorized as ***pre-Start G1***.

Nuclear mean pixel intensity of Whi5-WIQ-mCitrine was also used for assigning the timepoint at which cells enter telophase. The first frame of nuclear re-import of Whi5-WIQ-mCitrine in S/G2/M cells, was manually identified for individual cells in cell categories of interest. All frames including and after this frame were categorised as ***post-anaphasic***.

Background adjusted Htb2-mScarlet-I amount in the bud was used to assign the beginning of anaphase. The first frame showing increasing Htb2-mScarlet-I amount in the bud was manually identified for cells of interest. All frames after and including this frame and before the first telophase frame were categorised as ***anaphasic***.

Finally, all S/G2/M frames starting from and including bud emergence and going until the first anaphasic frame were assigned to the ***bud emergence to anaphase*** category, spanning S, G2, prophase and metaphase.

### Statistical Analyses

Except for the confidence interval comparison (Fig. 1G) and mixed ANOVA (Fig. 2B,D), all statistical analyses were done using independent two-tailed t-tests assuming unequal variances (Welch’s). p-values below 0.001 were denoted by ‘***’, p-values below 0.01 were denoted by ‘**’, p-values below 0.05 were denoted by ‘*’ and p-values > 0.05 were denoted by ‘ns’ for non-significant.

## Acknowledgements

We thank Benedikt Mairhörmann, Mardo Kõivomägi, Jennifer Ewald, Pascal Falter-Braun, Andreas Klingl and members of the Institute of Functional Epigenetics for helpful discussions. This work was funded by the Deutsche Forschungsgemeinschaft (DFG, German Research Foundation) through project 431480687, by the Human Frontier Science Program (career development award to K.M.S.), and the Helmholtz Gemeinschaft. Work in the RS lab was funded by the Deutsche Forschungsgemeinschaft (DFG, German Research Foundation) through SFB 1064 (Project-ID 213249687) and SFB 1309 (Project-ID 325871075) as well as the Helmholtz Gemeinschaft.

