## Supplementary Information for "Single-cell imaging reveals a key role of Bck2 in budding yeast cell size adaptation to nutrient challenges"

| Name | Genotype | Source | Figures used in |
|---|---|---|---|
| MMY116-2C | *MATα ADE2* | Lab Stock | "WT" in 1, 2, 3, 4, 5, 7, S2, S3, S4, S5 & S6 |
| YCY001-1 | *MATα ADE2 bck2Δ::CglaTRP1* | This study | "*bck2Δ*" in 1, 4, 5, 7, S2, S4, S5 & S6 |
| YCY002-1 | *MATα ADE2 ccr4Δ::CglaTRP1* | This study | "*ccr4Δ*" in 1, S2, S3 & S4 |
| YCY004-5 | *MATα ADE2 whi5Δ::KlacURA3* | This study | "*whi5Δ*" in 1, 4, 5 & 7 |
| YCY005-1 | *MATα ADE2 bck2Δ::CglaTRP1 whi5Δ::KlacURA3* | This study | "*whi5Δbck2Δ*" in 1, 2, 4, 5, 7, S4, S5 & S6 |
| YCY006-1 | *MATα ADE2 ccr4Δ::CglaTRP1 whi5Δ::KlacURA3* | This study | "*whi5Δccr4Δ*" in 1, S3 & S4 |
| YCY024-1 | *MATα ADE2 ura3::Act1promoter-WHI5WIQ-mCitrine-URA3 htb2::HTB2-mScarlet-I-natMX6* | This study | "WT" in 6 |
| YCY030-2 | *MATα ADE2 ura3::Act1promoter-WHI5WIQ-mCitrine-URA3 htb2::HTB2-mScarlet-I-natMX6 whi5Δ::CglaTRP1* | This study | "*whi5Δ*" in 6 |
| YCY037-1 | *MATα ADE2 ura3::Act1promoter-WHI5WIQ-mCitrine-URA3 htb2::HTB2-mScarlet-I-natMX6 bck2Δ::CglaLEU2* | This study | "*bck2Δ*" in 6 |
| YCY038-1 | *MATα ADE2 ura3::Act1promoter-WHI5WIQ-mCitrine-URA3 htb2::HTB2-mScarlet-I-natMX6 whi5Δ::CglaTRP1 bck2Δ::CglaLEU2* | This study | "*whi5Δbck2Δ*" in 6 |
| YCY021-2 | *MATα ADE2 cln3Δ::CglaLEU2* | This study | "*cln3Δ*" in 7 & S6 |
| YCY022-2 | *MATα ADE2 cln3Δ::CglaLEU2 whi5Δ::KlacURA3* | This study | "*whi5Δcln3Δ*" in 7 & S6 |
| YCY023-1 | *MATα ADE2 cln3Δ::CglaLEU2 whi5Δ::KlacURA3 bck2Δ::CglaTRP1* | This study | "*whi5Δbck2Δcln3Δ*" in 7 & S6 |

**Supplemental table 1:** Table describing the strains used in this study. All the strains used are haploid derivatives of W303, constructed using standard methods and verified by sequencing.

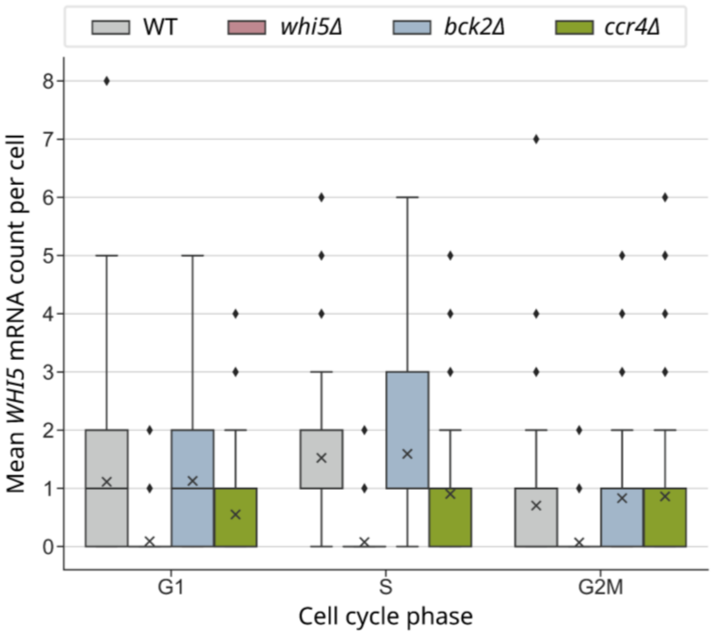


**Supplemental figure 1: Expression of *WHI5* peaks during S-phase.** Mean *WHI5* mRNA count per cell for cells in different cell cycle stages. Data is pooled from two independent smFISH experiments performed in SCD. G1-phase: $n_{WT}$ = 98, $n_{whi5\Delta}$ = 121, $n_{bck2\Delta}$ = 196, $n_{ccr4\Delta}$ = 127. S-phase: $n_{WT}$ = 172, $n_{whi5\Delta}$ = 182, $n_{bck2\Delta}$ = 211, $n_{ccr4\Delta}$ = 83. G2/M-phase: $n_{WT}$ = 125, $n_{whi5\Delta}$ = 143, $n_{bck2\Delta}$ = 178, $n_{ccr4\Delta}$ = 130.

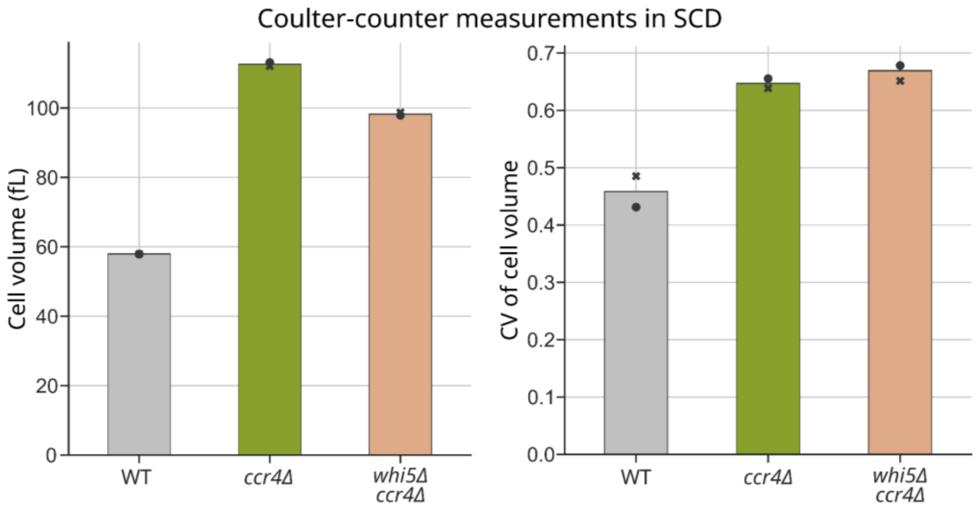


**Supplemental figure 2: *whi5Δccr4Δ* does not rescue the large size phenotype of *ccr4Δ.*** Coulter-counter measurements of cell volume (left) and CV of cell volume (right) are plotted for cells growing in SCD. Data is pooled from two independent experiments.

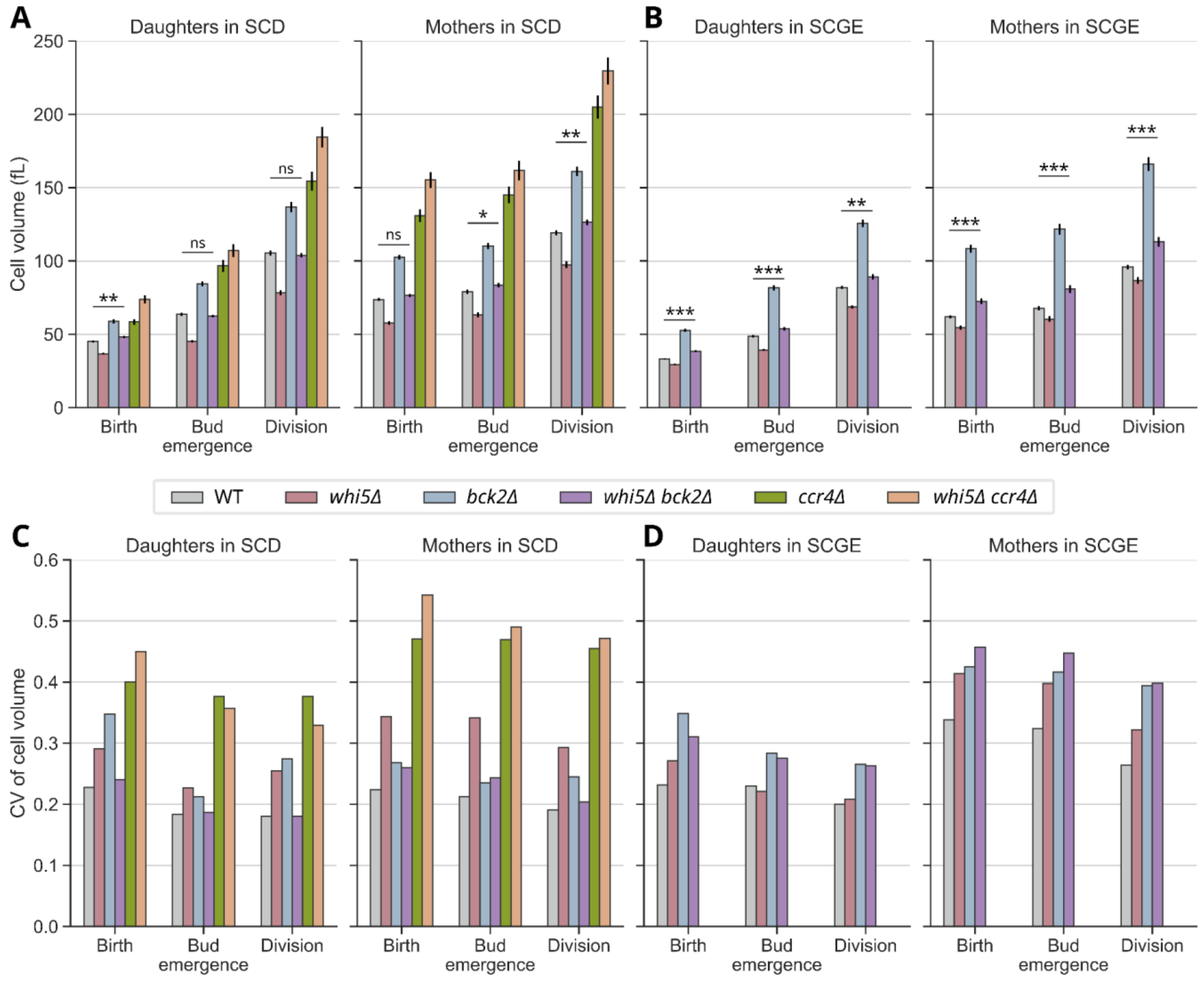


**Supplemental figure 3: Cell volume and CV of cell volume of mother and daughter cells at different cell cycle stages in two growth media.** Cell volume at birth, bud emergence and division is plotted for daughter cells and mother cells growing in SCD (**A**) and SCGE (**B**). CV of cell volume at birth, bud emergence and division is plotted for daughter and mother cells growing in SCD (**C**) and SCGE (**D**).

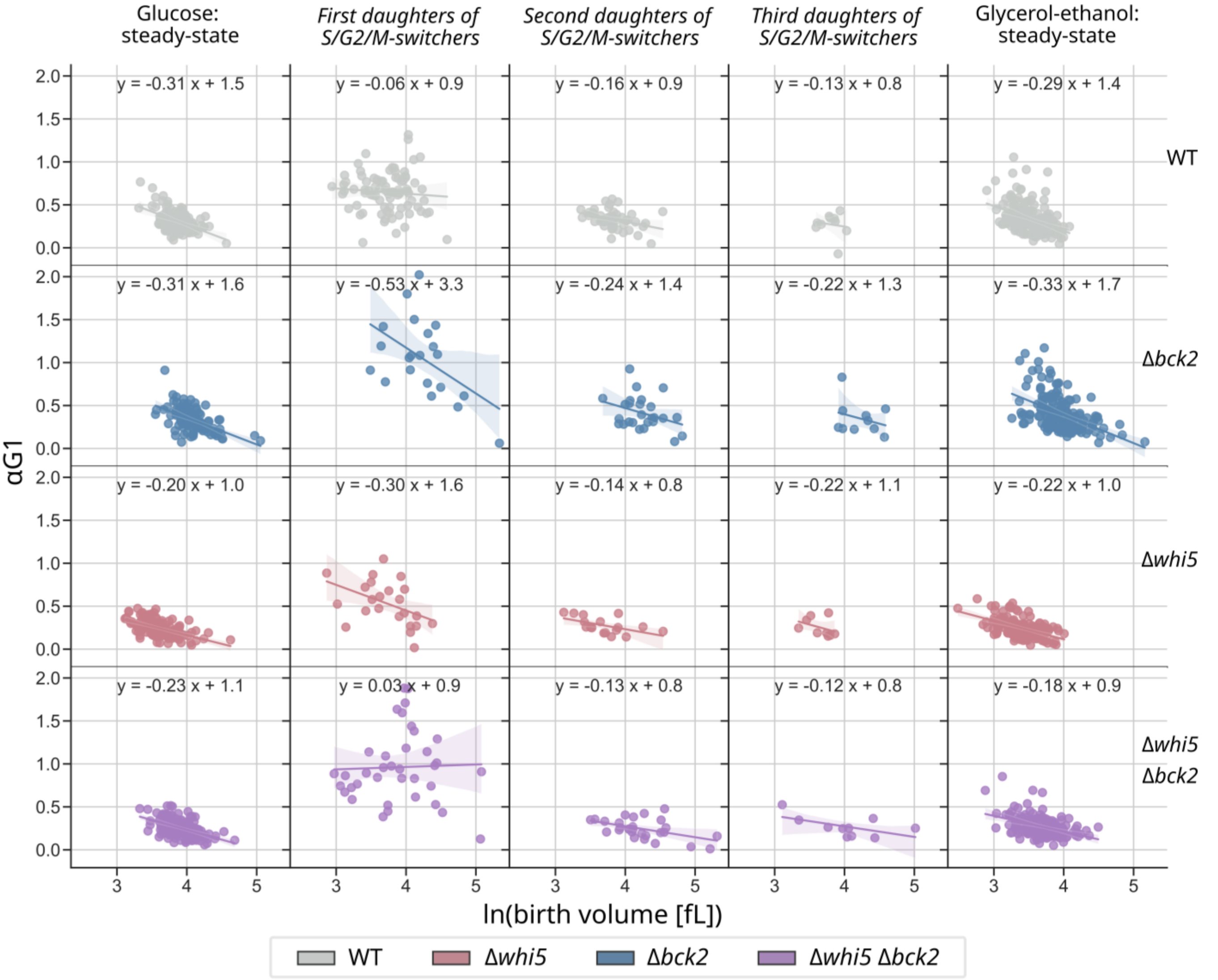


**Supplemental figure 4: Cell size homeostasis at the G1/S transition is weaker in the *daughters of S/G2/M switchers*.** This figure shows the strength of size homeostasis at the G1/S transition in different cell categories assessed as per the method described in [52]. The growth rate for exponentially growing cells, α, is the slope of the natural logarithm (in [fL]) of cell volume plotted against time. For each cell, G1 length is multiplied by α to obtain a growth rate adjusted G1-duration, αG1. αG1 is plotted against the natural logarithm of birth volume and a linear regression is performed. A negative slope of the resulting regression line indicates strong cell size homeostasis, whereas a slope closer to 0 indicates weaker cell size homeostasis. Shaded areas show 95% confidence intervals. While steady-state wild-type daughters (outermost left and outermost right panels) show a steeper negative slope and therefore stronger size homeostasis, the first round of daughters of *S/G2/M switchers* (WT, second panel) exhibits much weaker size homeostasis. The following rounds of wild-type daughters show slight recovery in size homeostasis strength. By contrast, the first round of daughters of *whi5Δ* and *bck2Δ S/G2/M-switchers*, but not those of *whi5Δbck2Δ S/G2/M switchers* show stronger size homeostasis. In steady-state, *whi5Δ* and *whi5Δbck2Δ* daughters have the lowest G1/S size control efficiency.

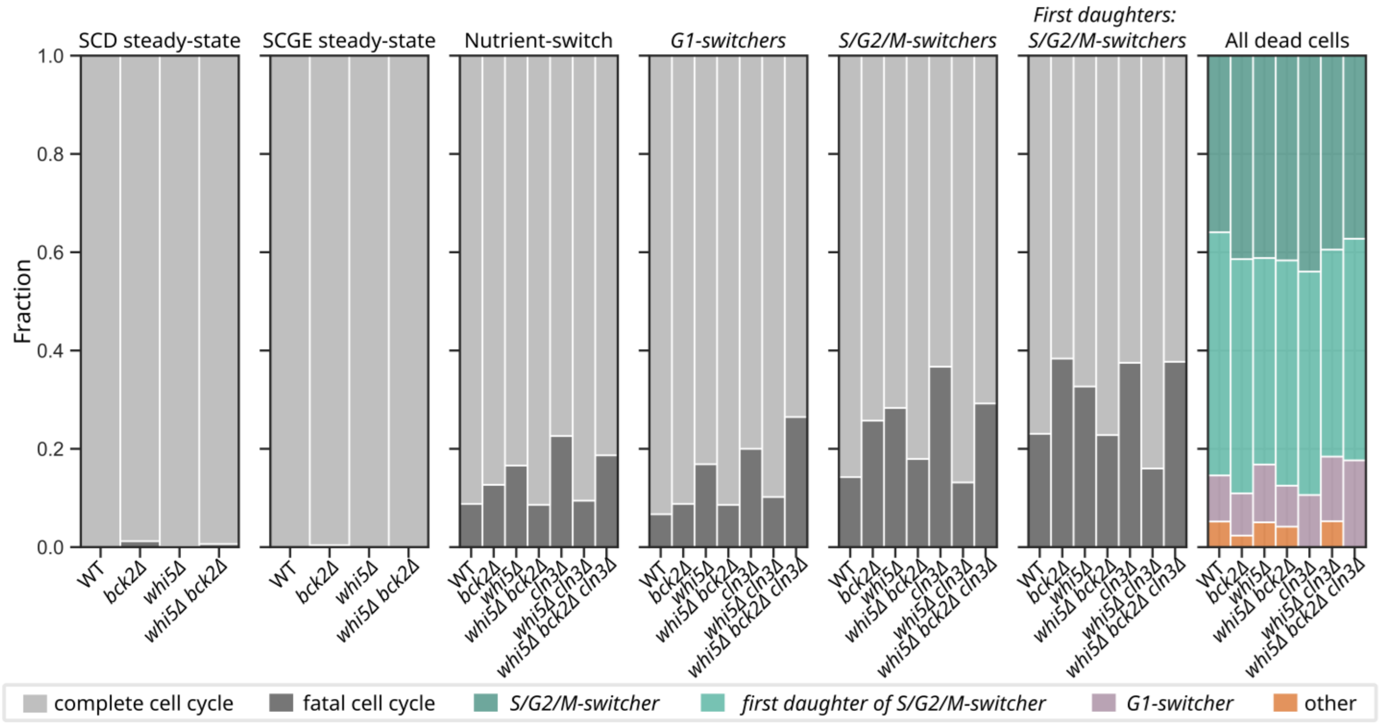


**Supplemental figure 5: *S/G2/M-switchers* and their *first daughters* make up the majority of dying cells.** For cells pooled from a minimum of two independent nutrient switch experiments, all complete cell cycles and cell cycles during which the cells died were identified. Of this total, the fraction of cell cycles during which cells died (fatal cell cycles, dark grey) and the fraction of cell cycles that were completed (complete cell cycle, light grey) are plotted in a stacked bar plot. The first two panels show the fraction of fatal cycles in steady-state in SCD and SCGE. The third panel shows the fraction of fatal cycles during or after the nutrient switch. In the nutrient switch, the single deletions, *whi5Δ*, *cln3Δ* and *bck2Δ*, and the triple deletion, *whi5Δbck2Δcln3Δ*, had increased fractions of fatal cell cycles as compared to WT. The double deletions, *whi5Δbck2Δ* and *whi5Δcln3Δ*, rescued survival. This may be linked to the double deletions being closer to WT size than the single deletions. Panels four, five and six show the fraction of fatal cycles if only cell populations of interest are considered: *G1-switchers*, *S/G2/M-switchers* and *first daughters of S/G2/M switchers*. In the last panel, all fatal cycles during and after the nutrient switch are pooled. The fraction of dead cells coming from different cell categories is plotted. *S/G2/M-switchers* and their *first daughters* make up the majority of dying cells.

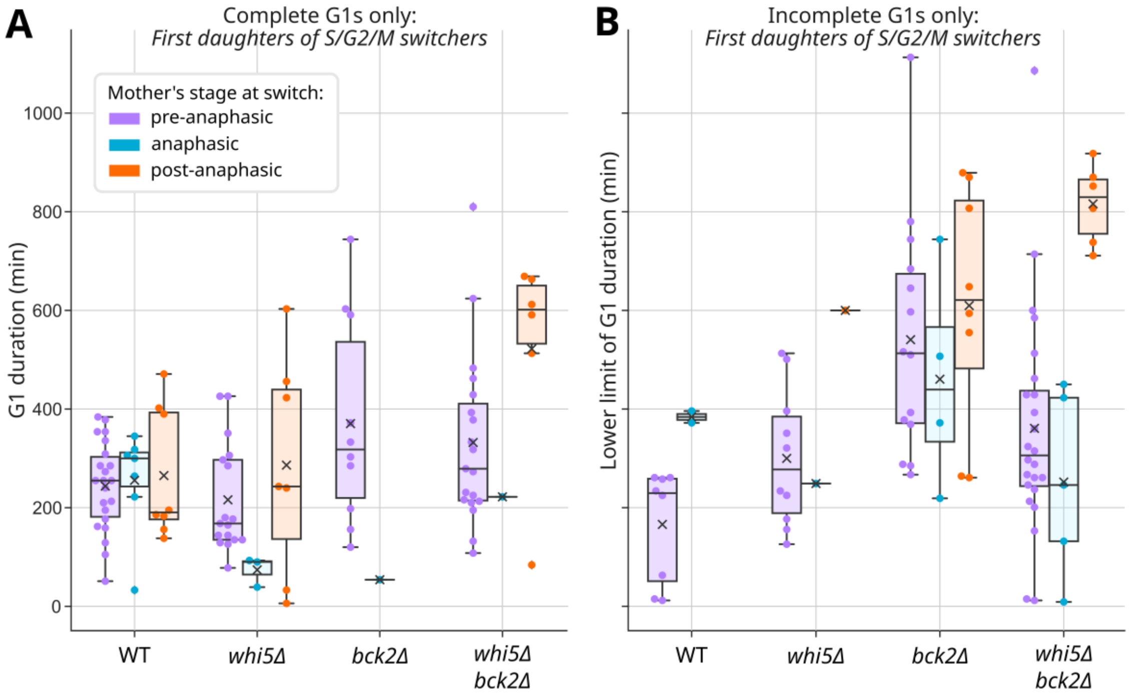


**Supplemental figure 6: In *bck2Δ* cells, the first daughters of mothers which faced the nutrient switch after anaphase have the strongest arrest phenotypes.** We categorised the first round of daughters of *S/G2/M-switchers* by whether the mother was pre-anaphasic (purple), anaphasic (blue) or post-anaphasic (orange) at the time of the nutrient switch and analysed the corresponding G1-lengths. The durations of complete and incomplete G1 phases of the first daughters of *S/G2/M-switchers* are plotted in (**A**) and (**B**) respectively. Incomplete G1 phases are G1 phases interrupted by the end of the experiment and not by death or exclusion of cells from the analysis. The y-axis of (**B**), therefore, is a lower limit of G1 duration. x symbol denotes the mean phase duration of the population. For *whi5Δbck2Δ* cells, the strongest arrest phenotype, *i.e.*, the longest G1 lengths, were observed for the group whose mothers were post-anaphasic at the time of nutrient switch (**A**). This was also reflected in the lower limit of G1 durations that we inferred for incomplete G1 phases (**B**). For *bck2Δ*, we did not observe any complete G1 phases for the group of daughters whose mothers were post-anaphasic at the time of nutrient switch (**A**) as all of these daughters stayed arrested in G1 until the end of the experiment (**B**).
